# High resolution profiling of cell cycle-dependent protein and phosphorylation abundance changes in non-transformed cells

**DOI:** 10.1101/2024.06.20.599917

**Authors:** Camilla Rega, Ifigenia Tsitsa, Theodoros I. Roumeliotis, Izabella Krystkowiak, Maria Portillo, Lu Yu, Julia Vorhauser, Jonathon Pines, Joerg Mansfeld, Jyoti Choudhary, Norman E. Davey

## Abstract

The cell cycle governs a precise series of molecular events, regulated by coordinated changes in protein and phosphorylation abundance, that culminates in the generation of two daughter cells. Here, we present a proteomic and phosphoproteomic analysis of the human cell cycle in hTERT-RPE-1 cells using deep quantitative mass spectrometry by isobaric labelling. Through analysing non-transformed cells, and improving the temporal resolution and coverage of key cell cycle regulators, we present a dataset of cell cycle-dependent protein and phosphorylation site oscillation that offers a foundational reference for investigating cell cycle regulation. These data reveal uncharacterised regulatory intricacies including proteins and phosphorylation sites exhibiting previously unreported cell cycle-dependent oscillation, and novel proteins targeted for degradation during mitotic exit. Integrated with complementary resources, our data link cycle-dependent abundance dynamics to functional changes and are accessible through the Cell Cycle database (CCdb), an interactive web-based resource for the cell cycle community.

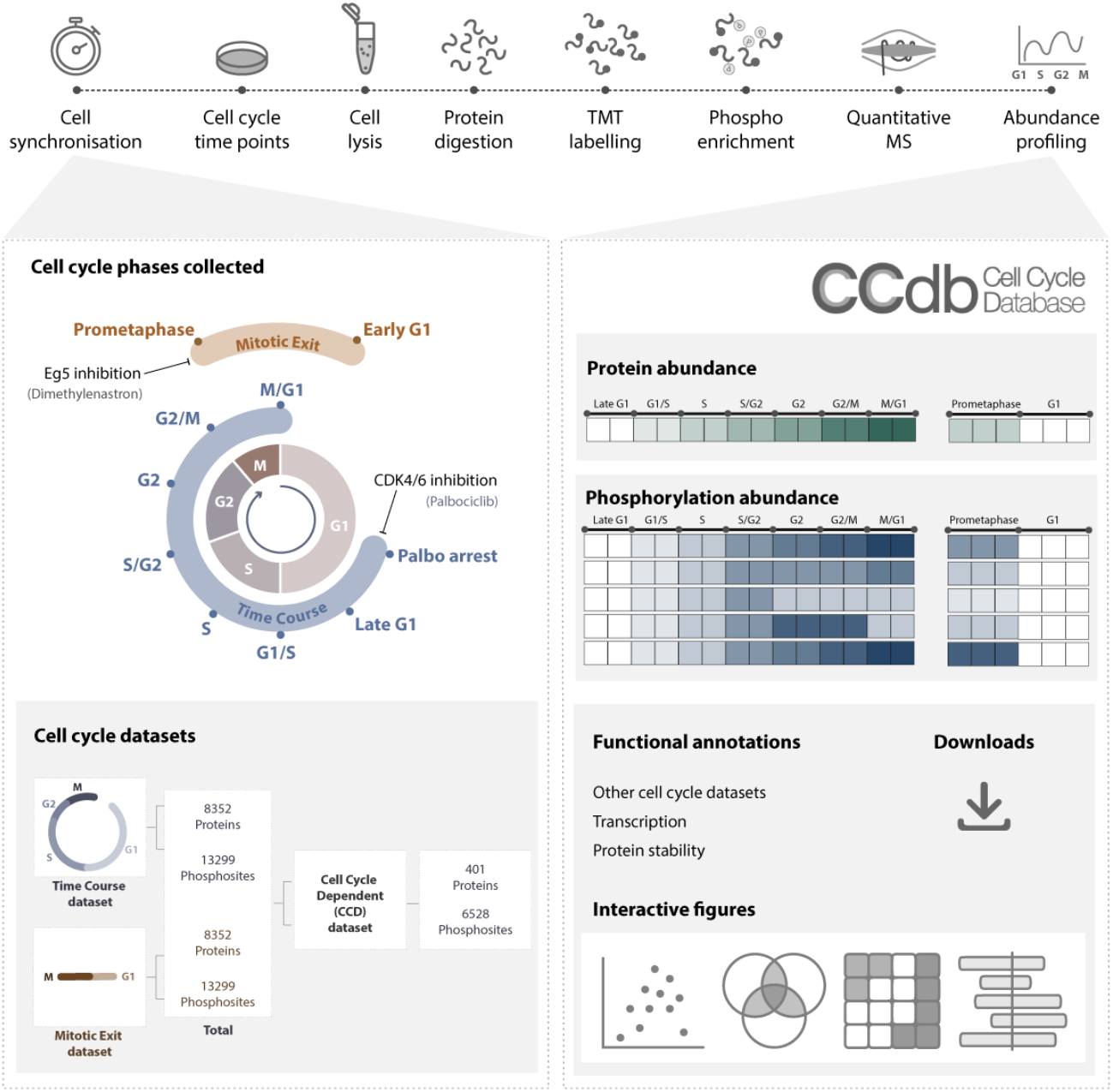

## Introduction

The eukaryotic cell cycle is a highly regulated sequence of events that culminate in cell division ^1^. It involves a complex network of regulatory proteins that ensure accurate progression through each phase of the cell cycle. Many of these proteins act in checkpoints, monitoring DNA replication and ensuring that the two daughter cells inherit an equal and identical set of chromosomes. Disruption of this intricate regulatory system can have catastrophic consequences, including uncontrolled cell proliferation and aneuploidy that are characteristics of diseases such as cancer. The cell cycle is regulated at multiple levels to ensure proper progression and control. One of the key mechanisms is the balance between transcription, translation and protein degradation ^2^, crucial for the temporal and spatial regulation of protein expression during specific stages of the cell cycle. In parallel, kinases and phosphatases work antagonistically with each other to dynamically control the phosphorylation state of individual proteins^3^.

Since the discovery of cyclins over 40 years ago ^4^, significant progress has been made in understanding the role of protein and phosphorylation oscillation in cell cycle progression. Decades of low-throughput experiments have uncovered waves of protein and phosphorylation abundance changes that robustly order the events required for cell division. However, a significant challenge lies in accessing scattered data and integrating the diverse findings from various studies across multiple experimental setups. Therefore, over the last years, several groups have investigated cell cycle regulation using Mass Spectrometry (MS) and provided valuable insights into the dynamic changes in protein and phosphorylation that occur throughout the cell cycle ^5–9^. While these datasets provide a robust foundation to investigate cell cycle dynamics, they have inherent limitations: the use of cancer cell lines, which may not accurately reflect the normal processes of the cell cycle, and the limited set of cell cycle phases collected through each study. Moreover, despite our growing understanding of protein dynamics in the cell cycle, there is still a significant gap in our knowledge when it comes to phosphorylation events, which are more technically challenging to detect and study in detail ^8^. Consequently, data from multiple datasets must be combined for a comprehensive understanding of cell cycle processes.

Several studies have used chemical synchronisation to collect cells in different cell cycle stages ^10–12^. However, these methods are often associated with a negative impact on the cell cycle and increased DNA damage ^10^. One way to address these challenges is through the use of fluorescent cell cycle reporters, such as the FUCCI system ^5,7^. Although these reporters overcome the need for synchronisation, data collection is often limited by the low number of cells processed at different stages of the cell cycle. Therefore, induction synchrony remains the most widely used approach due to its accessibility, efficiency and ability to preserve cell viability. The CDK4/6-targeting inhibitor Palbociclib has emerged as an effective and reversible tool to induce highly synchronised cell populations with minimal impact on the cell cycle by arresting cells at the natural restriction point ^12,13^. The high degree of synchrony achieved through palbociclib offers a robust framework for exploring the mechanisms of cell cycle regulation.

In this study, we present a high resolution quantitative proteomic and phosphoproteomic analysis of cell cycle progression that integrates data from other studies to provide a comprehensive resource. We use the hTERT-immortalised retinal pigment epithelial (RPE-1) cell line, which has a complete set of cell cycle regulators and is widely recognised as a model for studying cell cycle processes ^14^. We provide a map of the protein and phosphorylation dynamics through the cell cycle by performing deep MS analysis across seven distinct stages of the cell cycle after synchronisation with a palbociclib-induced reversible arrest. Additionally, we employed a prometaphase arrest and release into G1 phase to investigate events occurring during mitotic exit. These datasets include abundance profiles for thousands of proteins and phosphorylation sites, providing a valuable resource for studying the intricate regulatory mechanisms governing cell cycle progression. Through analysis of the resulting temporal profiles, we identified a high-confidence set of proteins exhibiting cell cycle-dependent abundance changes, establishing a gold standard reference for investigating cell cycle regulation. We validate novel degradation mechanisms for under characterised cell cycle proteins and show that a significant proportion of proteins with cell cycle-dependent oscillation patterns have no clear mechanism for targeted degradation. Finally, we characterise the context of cell cycle-dependent phosphorylation sites to predict protein interactions modulated in a cell cycle-dependent manner. The analysis is accompanied by the Cell Cycle database (CCdb), an online platform that enables easy access and exploration of the proteins or phosphorylation abundance profiles. The CCdb provides an in-depth resource for the cell cycle community to unravel the complexity of cell cycle regulation.

## Results

### High resolution time course proteomic and phosphoproteomic analysis of cell cycle progression in RPE-1 cells

To comprehensively capture the protein and phosphorylation dynamics during cell division in non-transformed human retinal pigment epithelial (RPE-1) cells, we coupled reversible cell cycle inhibition arrest with isobaric peptide labelling (TMT) quantitative Mass Spectrometry (MS). Cells were synchronised with palbociclib, a CDK4/6 inhibitor that arrests cells at the restriction point in late G1 and has recently been shown to generate a synchronous cell cycle upon inhibitor withdrawal ^12^. To generate a high temporal resolution cell cycle profile, we collected eight time points corresponding to different cell cycle phases: palbociclib arrest, late G1, G1/S, S, S/G2, G2, G2/M and M/G1 (Figure 1A, Supplementary Figure 1A). Two biological replicates were collected for each cell cycle phase. Cell synchronisation efficiency was assessed by monitoring DNA content changes over time using flow cytometry before performing MS analysis (Figure 1B). Synchronised cells were lysed, proteins digested with trypsin and peptides labelled with the TMTpro 16-plex reagents. For phosphoproteomics profiling, phosphopeptides were enriched using an Fe-NTA metal affinity resin and the flow-through fraction was used for proteomic analysis.

**Figure 1:**
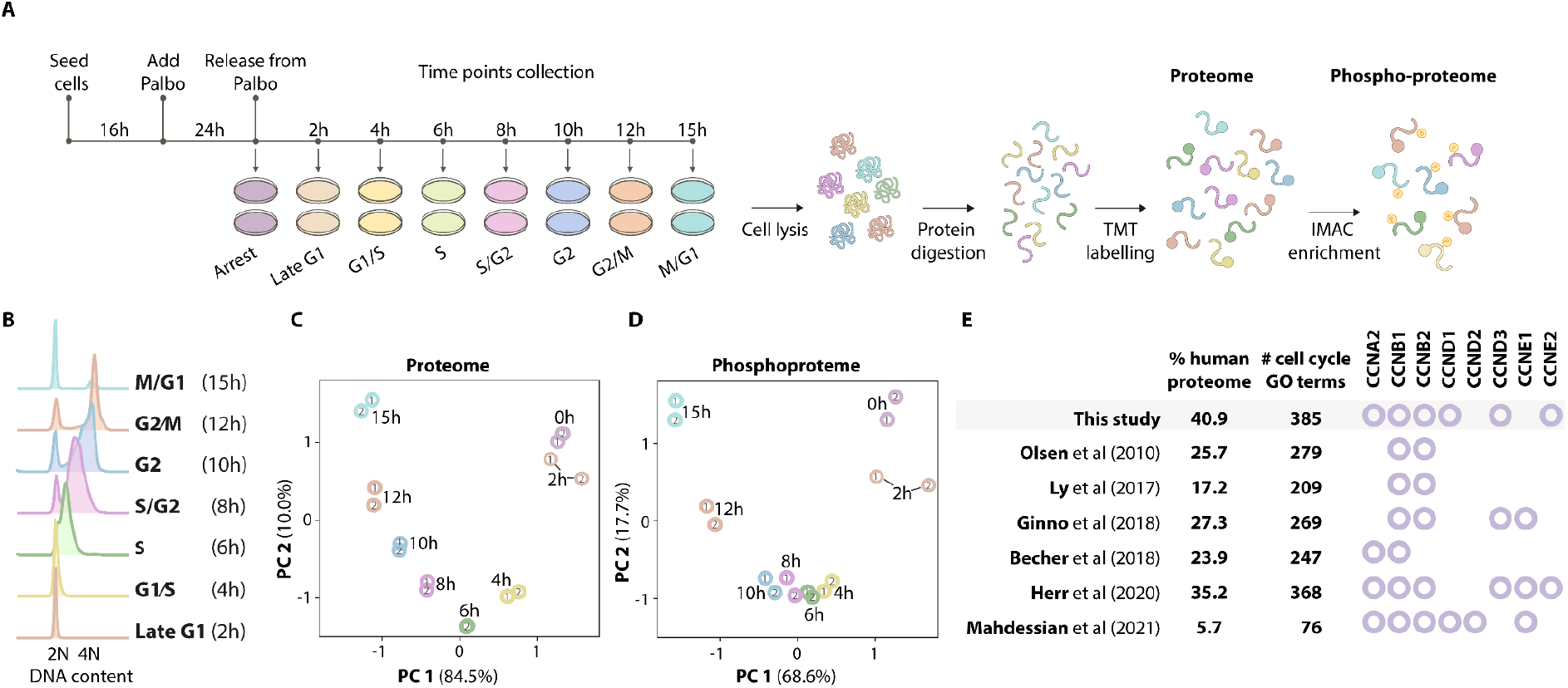
Quantitative proteomic and phosphoproteomic analysis of the cell cycle in RPE-1 cells. **(A)** General experiments workflow of cell synchronisation coupled to MS analysis. Cells were arrested in late G1 with palbociclib (Palbo) and time points corresponding to different cell cycle phases were collected upon release from the inhibitor. Sample aliquots were fixed and DNA was stained with propidium iodide for flow cytometry. For MS analysis, cells were lysed and protein digested with trypsin, followed by TMT labelling and phosphopeptide enrichment. **(B)** Flow cytometry analysis assessing DNA content at each time point. **(C)** Principal component analysis of the proteome. **(D)** Principal component analysis of the phosphoproteome. **(E)** Comparative analysis of our proteomics data with previously published cell cycle studies ^5–9,15^. For each dataset, the coverage of the human proteome (based on the 20,377 UniProt reviewed human proteins) and the number of proteins associated with the Gene Ontology (GO) term “mitotic cell cycle process” detected are listed. Cyclins identified in each study are shown with circles.

We quantified 8,352 proteins across the different cell cycle phases with high reproducibility within replicates (Figure 1C, Supplementary Figure 1B) covering a substantial proportion of the human proteome (nearly 41%), including 385 proteins associated with the Gene Ontology (GO) term “mitotic cell cycle process” (Figure 1E, Supplementary Table 1). Additionally, we detected 15,424 phosphorylation events, with 13,299 confidently assigned to a specific residue, also exhibiting high reproducibility within the replicates of each time point (Figure 1D). When comparing our dataset with other large-scale cell cycle studies using different cell lines (Olsen *et al* ^8^ (HeLa S3), Ly *et al* ^5^ (NB4), Ginno *et al* ^6^ (798G), Becher *et al* ^9^ (HeLa), Herr *et al* ^7^ (HeLa FUCCI), Mahdessian *et al* ^15^ (U2OS FUCCI) Supplementary Figure 1C), we not only identified a wider range of cell cycle regulators but also provided higher temporal resolution that enables an accurate representation of protein oscillation dynamics (Supplementary Figure 1D). This dataset, defined as the “Time Course dataset”, represents a comprehensive characterisation of the protein and phosphorylation dynamics during the cell cycle.

### Defining protein and phosphorylation oscillations during the cell cycle

Changes in protein and phosphorylation abundance determine the temporal progression of the cell cycle. Therefore, we first focused our analysis on defining oscillatory patterns using a curve-fitting model combined with ANOVA statistical analysis. The curve fitting model finds the curve that best fits a series of data points and provides a curve fold change score that describes to what extent protein and phosphorylation abundance changes over time. As shown in Figure 2A, we applied a two-step approach to define cell cycle-dependent oscillators in our proteomics and phosphoproteomics dataset. We first performed ANOVA analysis and found 459 proteins and 5,207 phosphorylation events showing clear changes between different cell cycle phases and positive correlation between replicates (q-values ≤ 0.01). Next, using the curve fitting model we calculated the curve fold change for each protein and phosphorylation site (Supplementary Table 2).

**Figure 2:**
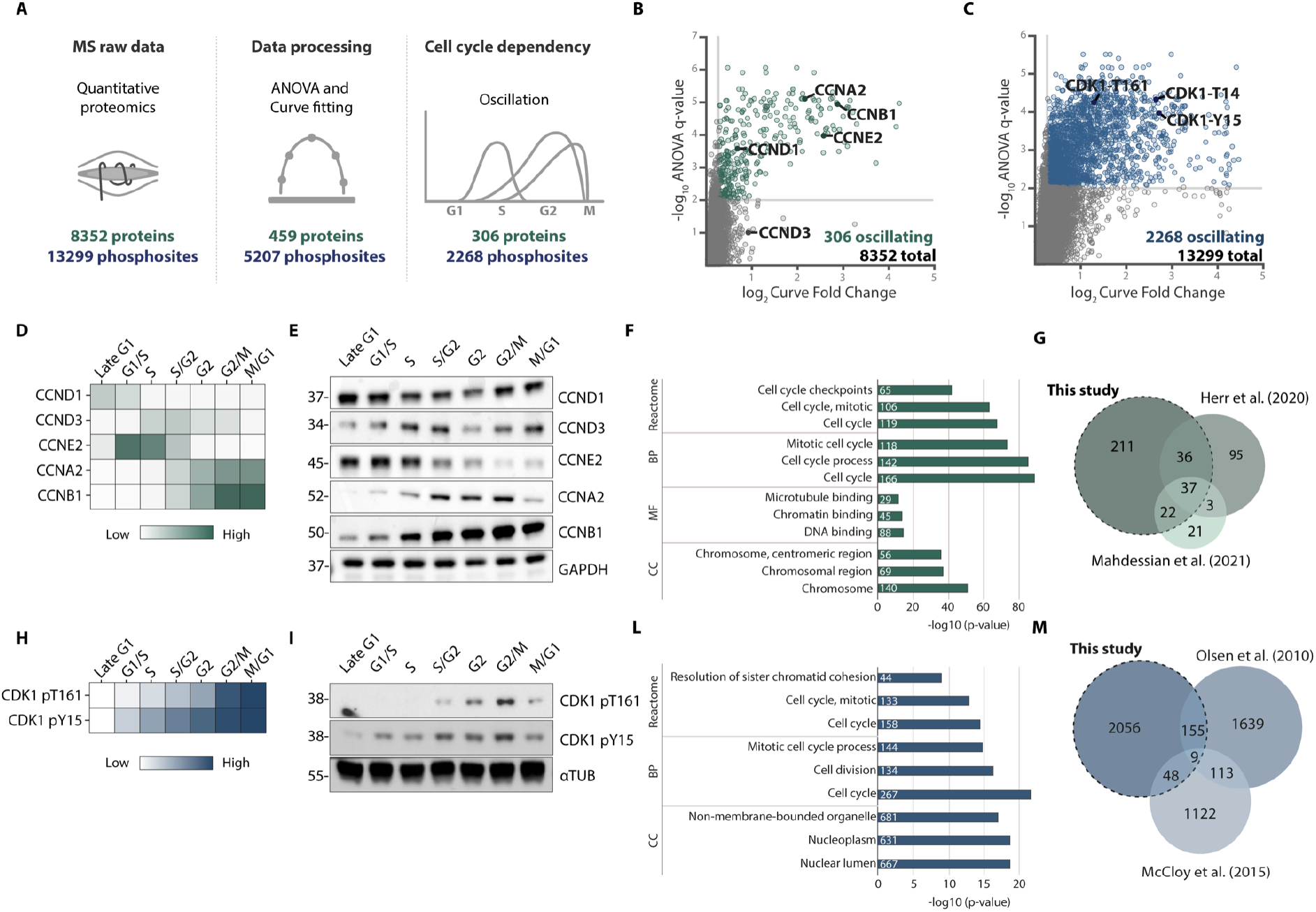
Defining oscillating protein and phosphorylation events during cell cycle progression. **(A)** Simplified data analysis workflow used to define oscillating proteins and phosphorylation sites. **(B)** Scatter plot summarising proteomics results shows the log_2_ curve fold change (oscillation score) and the -log10 q-values from the ANOVA test. Significantly oscillating proteins are shown in green and non-significant proteins are in grey. Dotted lines represent cut-offs used to define oscillating proteins (ANOVA q-values ≤ 0.01, curve fold change ≥ 1.2). Well-known cell cycle-regulated proteins are highlighted. **(C)** Scatter plot summarising phosphoproteomics results shows the fitted log_2_ curve fold change and the -log10 q-values from the ANOVA test. Significantly oscillating phosphorylation sites are shown in blue and non-significant phosphorylation sites are in grey. Dotted lines represent cut-offs used to define oscillating phosphorylation events (ANOVA q-values ≤ 0.01, curve fold change ≥ 1.2). Well-known cell cycle-regulated phosphorylation sites are highlighted. **(D)** Heatmap showing well-known cell cycle-regulated protein changes during the cell cycle detected by MS. Protein changes are coloured according to their abundance (log_2_ mean normalised values). **(E)** Western blot validation of the protein changes is shown in (D). **(F)** Overlap of our set of oscillating proteins with those detected in other studies. **(G)** Gene Ontology (GO) analysis of our set of oscillating proteins. The top three significantly enriched terms for each category (Reactome, Biological Processes, Molecular Function and Cellular Components) are shown. **(H)** Heatmap showing well-known cell cycle-regulated CDK1 phosphorylation changes during the cell cycle identified by MS. Phosphorylation changes are coloured according to their abundance (log_2_ mean normalised values). **(I)** Western blot validation of the phosphorylation changes shown in (H). **(L)** Overlap of our set of oscillating phosphorylation sites with those detected in other studies. **(M)** Gene Ontology (GO) analysis of oscillating phosphorylation sites. The top three significantly enriched terms for each category (Reactome, Biological Processes, Molecular Function and Cellular Components) are shown.

Based on the abundance pattern observed during the different stages of the cell cycle, proteins and phosphorylation events were categorised as oscillating and stable (Figure 2B and 2C). We found that 306 of 8,352 (3.7%) proteins and 2,268 of 13,299 (17%) phosphorylation events exhibited changes greater than 20% over the cell cycle (curve fold change ≥ 1.2, Supplementary Table 2). In contrast, 5,466 (65%) proteins and 593 (4.5%) phosphorylation events showed no significant difference within the different time points (ANOVA q-values ≥ 0.01, standard deviation ≤ 0.05). The cut-offs applied to define each category were defined based on GO term enrichment analysis to define biologically significant changes (Supplementary Table 3). Among the oscillating proteins, we identified several cyclins (i.e. CCND1, CCND3, CCNE2, CCNA2, CCNB1) (Figure 2D and 2E) and other well-known cell cycle regulators including CDK inhibitors (CDKN1A and CDKN1B), kinases (PLK1, AURKA and AURKB) and proteins regulating cell cycle checkpoints including WEE1, PKMYT1, CDC25A, CDC20 and PTTG1 (Supplementary Figure 2A and 2B) ^1^. Within the oscillating phosphorylation events, we observed numerous expected cell cycle-regulated phosphorylation events including the activating Thr161 and inhibitory Tyr15 phosphorylation sites of CDK1 increasing in G2/M ^16^ (Figure 2H and 2I). As expected, oscillating proteins and phosphorylation events were enriched for cell cycle-related Gene Ontology (GO) terms (Figure 2F and 2L, Supplementary Table 4).

Compared to two large-scale time course studies defining cell cycle oscillating proteins ^7,15^, our dataset revealed 211 proteins exhibiting previously uncharacterised changes in abundance during cell cycle progression (Figure 2G). The biological role of this set of novel oscillators was assessed using GO term analysis, showing a significant enrichment for protein associated with the “cell cycle” GO term (90 proteins, sig=1.5*x10*^−46^). Our data also included 95 proteins overlapping with one other cell cycle dataset ^7^, and 37 proteins were observed across all three studies ^7,15^ (Figure 2G). The largest overlap was observed with the Mahdessian *et al* oscillating set (71.1%, 59/83). The 24 proteins defined as oscillating only in the Mahdessian *et al* set included proteins that were not observed in our proteomic data (e.g., DBF4 and MYBL2), proteins that were oscillating without statistical significance (e.g., SGO1), or proteins where we observed a distinct paralogue as an oscillating protein (e.g., D- and E-type cyclins). There was a smaller overlap with the Herr *et al* oscillating set (42.7%, 73/171), however, we noted that the majority of proteins from this study were not observed in other datasets (55.6%) and these unique proteins were not enriched for “cell cycle” GO terms (16 proteins, sig=0.0023). The phosphoproteomic dataset revealed a greater amount of heterogeneity across various studies compared to proteomic data (Supplementary Figure 2C). Comparison with two large-scale studies defining cell cycle oscillating phosphorylation events ^8,17^ revealed only 212 oscillating phosphorylation events were also detected in these two datasets (Figure 2M).

### Protein and phosphorylation changes during mitotic exit

Our proteomics and phosphoproteomics analysis of cell cycle progression revealed that CDK4/6 inhibitor withdrawal provides a high degree of synchrony from late G1 to G2. However, because cells tend to differ in the rate at which they go through the cell cycle, synchrony was less robust at the entry to mitosis. As a result, investigating the changes occurring in mitosis with this synchronisation approach is challenging. Therefore, to improve our resolution from M to G1, we arrested cells in prometaphase using the Eg5 inhibitor Dimethylenastron (DMA) and subsequently released them to complete mitosis and progress into early G1. To prevent potential mitotic defects resulting from prolonged DMA treatment, cells were first synchronised in late G1 with palbociclib before DMA treatment in G2. In this analysis, we also included cells synchronised with serum deprivation to investigate the protein signatures of cells in G1 without the use of chemical agents. As shown in Figure 3A and Supplementary Figure 3A, this additional cell cycle dataset includes the following cell cycle phases: palbociclib arrest (late G1) - DMA arrest (prometaphase) - DMA release (early G1) - serum starvation arrest (G0) - serum starvation release (G1), defined as the “Arrest/Release dataset”. As in the Time Course dataset, synchronised cells were lysed, proteins digested with trypsin and peptides labelled with the TMTpro 16-plex reagents in three biological replicates. For phosphoproteomics profiling, phosphopeptides were enriched with an Fe-NTA metal affinity resin and the flow-through fraction was used for proteomics analysis. Before MS analysis, the efficiency of cell synchronisation was evaluated by monitoring changes in DNA content over time using flow cytometry (Supplementary Figure 3B).

**Figure 3:**
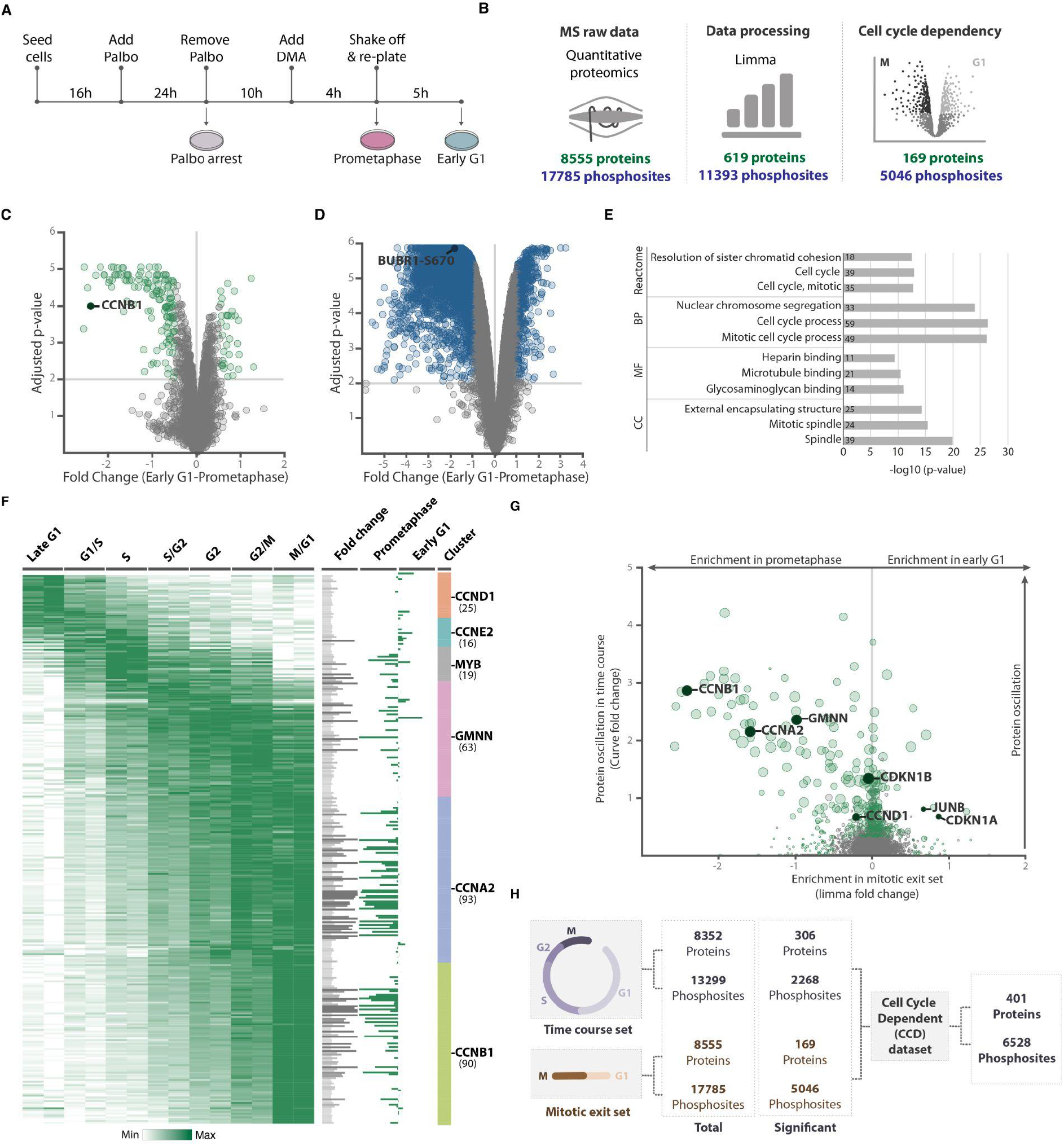
Protein and phosphorylation dynamics during mitotic exit. **(A)** Experimental workflow of cell synchronisation coupled to MS analysis. G2 phase cells pre-synchronised in G1 with palbociclib (Palbo) were incubated with DMA for 4h. Prometaphase-arrested cells were harvested by mitotic shake-off and released in fresh medium. 5h after release, early G1 cells were collected. For MS analysis, cells were lysed and protein digested with trypsin, followed by TMT labelling and phosphopeptides enrichment. **(B)** Simplified data analysis workflow used to define proteins and phosphorylation events significantly changing in prometaphase arrest and early G1. **(C)** Volcano plot showing limma adjusted p-values *versus* absolute limma log_2_ fold change of protein abundance in prometaphase arrest and early G1. Statistically significant proteins are shown in green, non-statistically significant proteins are shown in grey. Dotted lines represent cut-offs used to define oscillating proteins (limma adjusted p-value ≤ 0.001, log_2_ fold change ≥ 1.2). CCNB1 is significantly enriched in prometaphase arrest. **(D)** Volcano plot showing limma adjusted p-values versus absolute limma log_2_ fold change of phosphorylation sites abundance in prometaphase arrest and early G1. Statistically significant phosphorylation sites are shown in blue, non-statistically significant phosphorylation sites are shown in grey. Dotted lines represent cut-offs used to define oscillating phosphorylation events (limma adjusted p-value ≤ 0.001, log_2_ fold change ≥ 1.2). Phosphorylation at Ser670 of BUBR1 is significantly enriched in prometaphase arrest. **(E)** Gene Ontology (GO) analysis of proteins significantly changing in the Mitotic Exit dataset. The top three significantly enriched terms for each category (Reactome, Biological Processes, Molecular Function and Cellular Components) are shown. **(F)** Heatmap showing oscillating proteins detected in the Time Course dataset clustered based on the abundance profile of known cell cycle markers using T-distributed stochastic neighbour embedding. Protein changes are coloured according to their abundance (mean-max normalised values). Bar plot on the right shows curve fold change (oscillation score) in grey and protein differentially enriched in prometaphase or early G1 in the Mitotic Exit dataset in green. Clusters are grouped by colour and their respective reference protein is reported on the right. **(G)** Scatterplot comparing protein changes in the Mitotic Exit (fold change) and Time Course datasets (curve fold change). Statistically significant proteins are shown in green, non-statistically significant proteins are shown in grey. Point size denotes limma adjusted p-values from the Mitotic Exit dataset. Well-known cell cycle markers are highlighted. **(H)** Overview of the datasets presented in this study. The Time Course dataset covers seven cell cycle stages from late G1 to M/G1. The Mitotic Exit dataset involves prometaphase arrest and early G1. The Cell Cycle-Dependent (CCD) set combines proteins and phosphorylation sites that exhibit changes in the Time Course and Mitotic Exit datasets. The total number of proteins and phosphorylation sites in each dataset is shown.

TMT-based analysis revealed 8,555 proteins and 17,785 phosphorylation sites, quantified with a strong correlation between the three replicates (Supplementary Figure 3C, Supplementary Table 5). Comparison between the G0 and G1 phases from Serum Starvation arrest and release, in the “Serum Starvation dataset”, revealed minimal changes. However, in line with previous studies ^18^, we observed a significant depletion in CCND1 levels and an enrichment in CDKN1B expression in the serum starvation arrest compared to the release samples (Supplementary Figure 3D). There were also no major changes between late G1 from the Time Course dataset and G1 samples from the Serum Starvation dataset (Supplementary Figure 3E). As expected, pronounced changes were observed when comparing the prometaphase arrest to the early G1 phase in the DMA Arrest/Release experiments, defined as the “Mitotic Exit dataset”. Consistent with the dynamic nature of regulatory processes mediated by phosphorylation during mitosis, we found a significant difference in the overall levels of phosphorylation between prometaphase and early G1 (Supplementary Figure 3F). To address this discrepancy, we corrected the total intensity of phosphopeptides in the DMA arrest samples based on the stable phosphorylation sites detected in the Time Course set (Supplementary Figure 3G, see methods). Next, to identify proteins and phosphorylation events strongly modulated at the M to G1 transition we employed limma statistical analysis and defined cut-offs based on the enrichment of “cell cycle” GO term (Figure 3B, Supplementary Table 3). Using this approach, we identified 169 proteins and 5,046 phosphorylation sites significantly enriched in prometaphase or early G1 phase (limma adjusted p-values ≤ 0.001, absolute limma fold change ≥ 0.5, Figure 3B). Among the significantly changing proteins, we detected numerous known Anaphase-promoting complex/cyclosome (APC/C) substrates. For example, CCNB1 was significantly enriched in prometaphase arrest compared to early G1, consistent with its metaphase degradation required to exit from mitosis ^19^ (Figure 3C). We also detected significant changes for several well-characterised cell cycle-dependent phosphorylation sites, including the enrichment of BUB1 phosphorylation at Ser670 during prometaphase (Figure 3D). BUB1 is known to be phosphorylated at kinetochores at the beginning of mitosis and subsequently dephosphorylated before anaphase to facilitate proper chromosome alignment and error correction ^20^. Proteins significantly changing in the Mitotic Exit set showed significant enrichment of GO terms related to cell cycle processes, chromosome segregation and spindle organisation (Figure 3E, Supplementary Table 6).

### Comparing time course and mitotic exit datasets

Given that the Mitotic Exit set introduced an additional layer of specificity to the Time Course dataset, we anticipated a substantial overlap, especially among proteins peaking in mitosis. Therefore, we first clustered oscillating proteins identified in the Time Course dataset based on their abundance profile similarity with known cell cycle markers (Figure 3F). Reference proteins used in this analysis were CCND1, CCNE2, MYB, GMNN, CCNA2 and CCNB1. Next, we annotated proteins significantly enriched in the Mitotic Exit dataset for each protein found in the clusters. Overall, the largest clusters (i.e. CCNA2 and CCNB1) comprised proteins peaking in mitosis in both proteomics datasets. In contrast, clusters corresponding to proteins peaking in late G1, G1/S and S were under-represented in our analysis, suggesting that protein oscillation changes are modest in the early stages of the cell cycle. Furthermore, protein abundance over time clustered into different temporal “waves” of oscillation. Although the majority of the proteins peaked at the end of the cell cycle, specific protein clusters started accumulating at different cell cycle phases. For instance, proteins found to cluster with GMNN increased from S phase, while protein clustering with CCNA2 started to rise from S/G2, despite both clusters reaching their peak in G2/M. GO term enrichment analysis for each cluster revealed an overall enrichment in the expected cell cycle phase-specific terms (Supplementary Table 7). For example, the CCNE2 cluster was enriched for “cell cycle G1/S phase transition”, the MYB and GMNN cluster for “DNA replication”, the CCNA2 cluster for “chromosome organisation”, and the CCNB1 cluster for “chromosome segregation”. Moreover, the CCNA2 and CCNB1 clusters showed enrichment in mitotic localisation GO terms (kinetochore, spindle, centriole). This cluster analysis also revealed a strong correlation between proteins with high curve fold change scores (oscillation) in the Time Course dataset and fold change in the Mitotic Exit dataset (Figure 3G), particularly for proteins with increased abundance in mitosis (e.g., CCNA2, CCNB1 and GMNN). However, we also observed overlap between proteins peaking in late G1 in the Time Course and early G1 from the Mitotic Exit dataset (e.g. CDKN1A, CDKN1B, JUNB and CCND1). Similarly to the protein abundance, when comparing the Time Course with the Mitotic Exit dataset a significant overlap was observed at both the protein and phosphorylation level (Supplementary Figure 3H). Therefore, we combined the Time Course with the Mitotic Exit dataset to define a single dataset of Cell Cycle-Dependent (CCD) proteins and phosphorylation sites (Figure 3H) that we functionally investigate in the next sections. Moreover, the CCD proteins, clustered based on their temporal oscillation profile, discussed in this section can serve as a valuable reference set for investigating and comparing oscillation patterns, providing invaluable insights into cell cycle signatures (Supplementary Table 8).

### Biological role of cell cycle-dependent proteins

We further characterised the CCD protein set by integrating data on biological function, localisation, protein complexes and gene essentiality to investigate the functional role of protein oscillation during the cell cycle. As expected, given the observed enrichments of the Time Course and Mitotic Exit protein datasets, CCD proteins were highly enriched in cell cycle annotations, cell cycle-specific cellular structures (e.g., the kinetochore) and cell cycle-related pathways (Figure 4A, full GO terms list Supplementary Table 9). Next, we investigated the oscillation pattern of proteins within the same complex during cell cycle progression. We identified proteins belonging to the same complex using the Complex Portal and the CORUM databases ^21,22^ or based on their localisation using GO Cellular Component annotations (Supplementary Table 10). We assessed the variability of protein oscillation within the complexes (Figure 4B), revealing three main groups based on their oscillation patterns ^23^.

**Figure 4:**
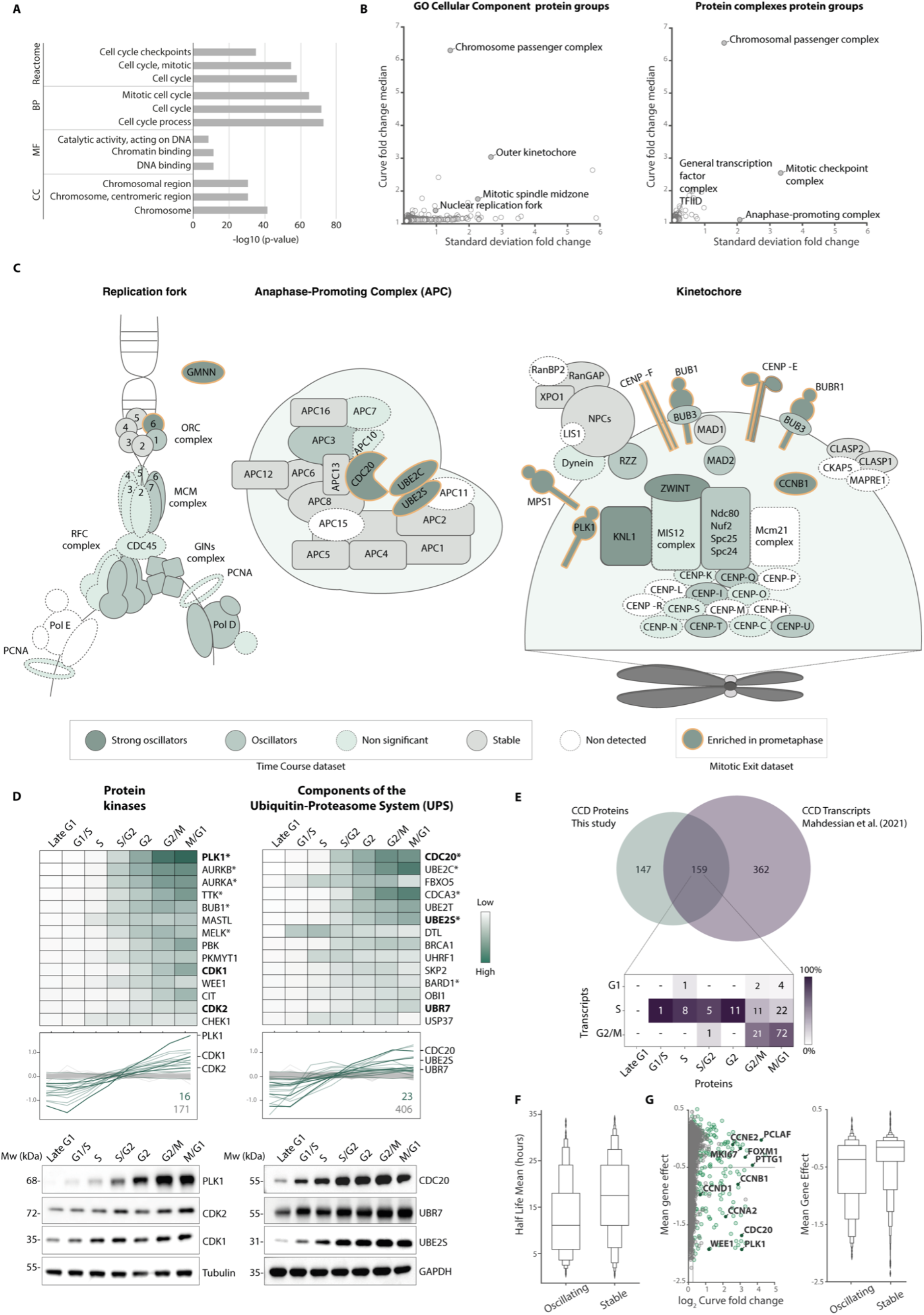
Functional significance of Cell Cycle-Dependent (CCD) proteins. **(A)** Gene Ontology (GO) analysis of CCD proteins. The top three significantly enriched terms for each category (Reactome, Biological Processes, Molecular Function and Cellular Components) are shown. **(B)** The oscillation of CCD proteins within the same cellular localisation or complex. Scatter plot showing the curve fold change median (global oscillation of proteins within each complex) and standard deviation (global variance in oscillation within each protein group) for protein sets grouped based on Gene Ontology-defined protein localisation (left panel) or proteins within the same complex obtained from CORUM and Complex Portal complexes ^21,22^ (right panel). **(C)** Schematic of CCD proteins localised at the replication fork, Anaphase Promoting Complex/Cyclosome and kinetochore. Protein colour indicates protein abundance detected in the Time Course dataset (Dark green: curve fold change > 2.5; Green: curve fold change > 1.2; Grey: curve fold change < 1.2 and standard deviation ≤ 0.05; White: not detected). Dotted borders indicate non statistically significant proteins. Yellow borders indicate proteins enriched in prometaphase in the Mitotic Exit dataset. **(D)** Heatmaps of the kinases and components of the ubiquitin-proteasome system (UPS) detected in the Time Course dataset (top panel). Protein changes are coloured according to their abundance (log_2_ mean normalised values). In the middle panel, profile plots showing the most significantly oscillating proteins. Number of oscillating (green) and stable (grey) proteins are shown and proteins validated by Western blot (bottom panel) are in bold. Proteins significantly enriched in the Mitotic Exit dataset are marked with an asterisk. **(E)** Venn diagram showing the overlap between oscillating proteins detected in the Time Course proteomics dataset and oscillating transcripts obtained from Mahdessian *et al.,* 2021. The heatmap shows the correlation between oscillating proteins and corresponding transcripts in each cell cycle phase. **(F)** Boxen plot comparing oscillating and stable proteins detected in the Time Course proteomics dataset with protein half-life values obtained from a cycloheximide chase proteomics dataset ^27^. **(G)** Essentiality of oscillating proteins detected in the Time Course proteomics dataset. Scatter plot comparing the curve fold change (oscillation score) and mean gene effect obtained from the Cancer Dependency Map resource ^28^. A lower score indicates a higher probability of gene essentiality, with a score of zero indicating that the gene is not required for cell survival, and a score of −0.5 suggesting essentiality in most cell lines. CCD proteins are shown in green, and non-CCD proteins are shown in grey. Well-known cell cycle markers are highlighted. Boxen plot shows a significant association between protein oscillation and lower mean gene effect scores (Mann-Whitney p-value: 2.76×10^−17^).

The first group consisted of stable complexes with low variability, characterised by consistent protein levels throughout the cell cycle. One example is the general transcription factor complex TFIID, which functions as a general transcription factor for transcription initiation by RNA polymerase II. The second group comprised complexes that exhibited high oscillation with low variability. In these complexes each protein is co-oscillating, for example, the chromosomal passenger complex where each subunit displayed oscillation behaviour through the cell cycle, reflecting its essential role in regulating chromosome segregation and cytokinesis. The third group included complexes with high variability where only specific subunits are cell cycle regulated while the remaining subunits are stable. For example, the median oscillation of proteins localised at the replication fork was relatively small, however, specific components, including ORC6 and GMNN, exhibited strong oscillatory behaviour (Figure 4B and 4C). Conversely, other ORC subunits were stable throughout the cell cycle (Supplementary Figure 4A). Similarly, the scaffolding proteins within the APC/C complex remained constant, while several regulatory subunits oscillated (Figure 4C and Supplementary Figure 4A). The APC/C activator CDC20, which plays a crucial role in promoting the degradation of various mitotic regulators, exhibited a strong oscillatory behaviour within the complex together with the ubiquitin-conjugating enzymes UBE2C and UBE2S. Within the kinetochore, a multitude of components were strongly oscillating and peaking in mitosis, including CDC20, MPS1, PLK1, BUB3, BUB1, BUBR1, CENPE, SWINT, CENPF, and KNL1 (Figure 4C and Supplementary Figure 4A). These components play critical roles in regulating diverse aspects of kinetochore function, including spindle checkpoint control, microtubule interactions, and chromosome attachment. In contrast, the inner kinetochore components, which provide structural integrity and support proper kinetochore function, largely remained stable throughout the cell cycle. Notably, our dataset revealed a distinct oscillation profile for CENPQ, peaking in G2, while its phosphorylation on the Ser50 was found to peak specifically during mitosis (Supplementary Figure 4B). This suggests that CENPQ plays an important role in the recruitment of key kinetochore components in agreement with previous findings ^24^.

To explore the cell cycle-dependent oscillation of regulatory proteins, we focused on specific classes of proteins known to play key roles in cell cycle progression and investigated their oscillation dynamics using the Time Course proteomics data. We classified the CCD proteins into three distinct regulatory classes: protein kinases, components of the ubiquitin-proteasome system (UPS) and transcription factors (TFs). We found that only a subset of these regulatory proteins exhibited oscillatory patterns during the cell cycle. Out of 187 kinases detected, only 16 showed significant oscillation, including CDK1, CDK2 and PLK1 (Figure 4D). Notably, eight of these protein kinases had not been previously reported to oscillate in other studies ^7,15^. Interestingly, we observed a continuous increase in the levels of CDKs during S phase, persisting through the G2/M phase, which contrasts with the conventional understanding that CDKs remain stable during the cell cycle and their activity is primarily regulated by cyclins and regulatory phosphorylation. This effect was particularly pronounced for CDK1, in agreement with previous studies which showed that the transcription of CDK1 is cell cycle regulated ^15,25,26^. Similarly to the kinases, among the 429 observed components of the UPS, 24 revealed significant oscillation (Figure 4D). These included several APC/C components, such as the previously mentioned CDC20 and UBE2S. Immunoblot analysis of synchronised cells confirmed the temporal changes detected by MS and validated the oscillation of the ubiquitin ligase UBR7, not previously described to be cell cycle regulated (Figure 4D). In contrast to the kinases and UPS components, the oscillation of TFs was generally less pronounced (Supplementary Figure 4C). However, a handful of TFs (14), including FOXM1, MYB, and JUNB, exhibited high levels of oscillation, consistent with their established role in cell cycle regulation and transcriptional control. Notably, among the 14 TFs detected in our study, only four of these were annotated with the “cell cycle” GO term, and none of them had been previously reported to oscillate in other studies ^7,15^.

To further investigate the link between transcription and protein oscillation during the cell cycle, we integrated our Time Course proteomics data with mRNA profiles from the Human Protein Atlas ^15^ (Supplementary Table 11). We found that approximately half of the oscillating proteins (159 out of 306, Supplementary Figure 5) also displayed oscillations on the transcript level, suggesting there is a strong link with transcriptional regulation (Figure 4E). Conversely, the remaining oscillating proteins (147 out of 306, Supplementary Figure 5) were found to have stable transcripts, suggesting that their regulation occurs primarily at post-transcriptional level and protein stability plays a significant role in determining their cell cycle-dependent expression dynamics. We further integrated protein half-life data obtained from cycloheximide chase experiments ^27^ to gain insights into the contribution of protein degradation to the oscillatory behaviour during the cell cycle (Supplementary Table 8). We anticipated that oscillating proteins would exhibit higher turnover rates compared to non-oscillating proteins. Consistent with our expectations, this analysis revealed a significant difference (Mann-Whitney p-value: 1.6*x10*^−14^) in protein half-lives between oscillating (mean of 12.1 hours, 25.1% less than 5 hours) and non-oscillating proteins (mean of 16.8 hours, 10.8% less than 5 hours)(Figure 4F). This observation underscores the impact of protein degradation in shaping the oscillatory behaviour of proteins during the cell cycle.

To explore the essentiality of cell cycle-regulated proteins in cell survival, we investigated the connection between CCD proteins and gene essentiality using the Mean Gene Effect scores obtained from CRISPR gene knockout and cell viability assays available in the Cancer Dependency Map resource ^28^ (Supplementary Table 8). The Mean Gene Effect score represents the likelihood that a particular gene is essential across a diverse set of cell lines. In agreement with previous findings ^7^, we observed a significant association between protein oscillation and lower Mean Gene Effect scores (Figure 4G). Among the CCD proteins detected in this study, we found several essential proteins, including CCND1, CCNB1, CCNA2, PLK1, WEE1 and CDC20. Our dataset also revealed some essential proteins not previously reported to oscillate in other studies ^7,15^. These include several components of the replication fork (e.g. GINS1, GINS2, MCM7, MCM6, ORC6, POLD1, POLD3, POLE, POLA2, RFC2 and RFC5). Interestingly, there were also CCD proteins that were not classified as essential for cell viability, such as CCNE2, PCLAF, FOXM1, and MKI67 (Figure 4G). These findings raise intriguing questions regarding redundancy in protein function and why proteins that have no effect on cell proliferation when knocked out are highly regulated through the cell cycle.

### Phosphorylation dynamics driving cell cycle progression

Protein phosphorylation is widely recognised as one of the major mechanisms regulating cell cycle progression ^29^. In this study, we identified a total of 22,305 phosphorylation events in the Time Course and Mitotic Exit datasets. Of these, 6,528 phosphorylation sites mapping to 4,693 proteins exhibited CCD behaviour (Figure 5A). This set of phosphorylation events was predominantly on serines (78.6%) and threonines (21.2%) with only a minor fraction occurring on tyrosines (0.1%) (Supplementary Figure 6A). As expected, both CCD and non-CCD phosphorylation sites were enriched within disordered and accessible regions of the proteins (Figure 5B, Supplementary Figure 6B). Since functionally important phosphorylation sites have been shown to be more likely to be conserved across different species ^30^, CCD phosphorylation sites were annotated with conservation scores quantifying the degree to which each site is conserved relative to its surrounding regions using the PepTools peptide annotation tool ^31^. Interestingly, we found that non-CCD phosphorylation sites were on average more conserved than CCD phosphorylation sites (Figure 5C). However, many CCD phosphorylations were highly conserved including phosphorylation events implicated in important cell cycle regulation mechanisms such as the phosphorylation of the Thr395 of CCNE1, which is known to regulate CCNE1 ubiquitination by the Fbw7 regulatory subunit-containing Skp, Cullin, F-box (SCF) E3 Ubiquitin ligase complex and subsequent targeting for degradation ^32^, and the phosphorylation of the kinesin-like protein KIF11 at Thr927, required for spindle association ^33^. We further investigated the taxonomic range of the CCD phosphorylation set and found that 72.4% of the phosphorylation sites were conserved to mouse (*M. musculus*), 44.5% to fish (*D. rerio*), 15.6% to fly (*D. melanogaster*), 3.3% to budding yeast (*S. cerevisiae*) and 5.5% to plant (*A. thaliana melanogaster*) (Figure 2D). Finally, to further investigate the temporal dynamics of phosphorylation regulation during cell cycle progression, we predicted the specific protein kinases responsible for CCD phosphorylation events. Using consensus motifs encoding the specificity of CDK, AURK, PLK and PIKK kinases, we classified all phosphorylation sites observed in the study. Notably, we observed a mirroring pattern of oscillating proteins and phosphosites for their respective kinases (Figure 5E). Substrates predicted for CDK exhibited the strongest oscillation, with enrichment beginning in G1/S and peaking late in the cell cycle, reflecting the known dynamics of CDK activity. PLK peaks late in the cell cycle, however, PLK sites accumulate after CDK sites in agreement with kinase activity organisation in sequential waves during the cell cycle. Furthermore, we observed an increase of PIKK substrates during the S phase in line with their known role in the DNA damage response ^34^ despite observing a stable abundance through the cell cycle for ATM and ATR.

**Figure 5:**
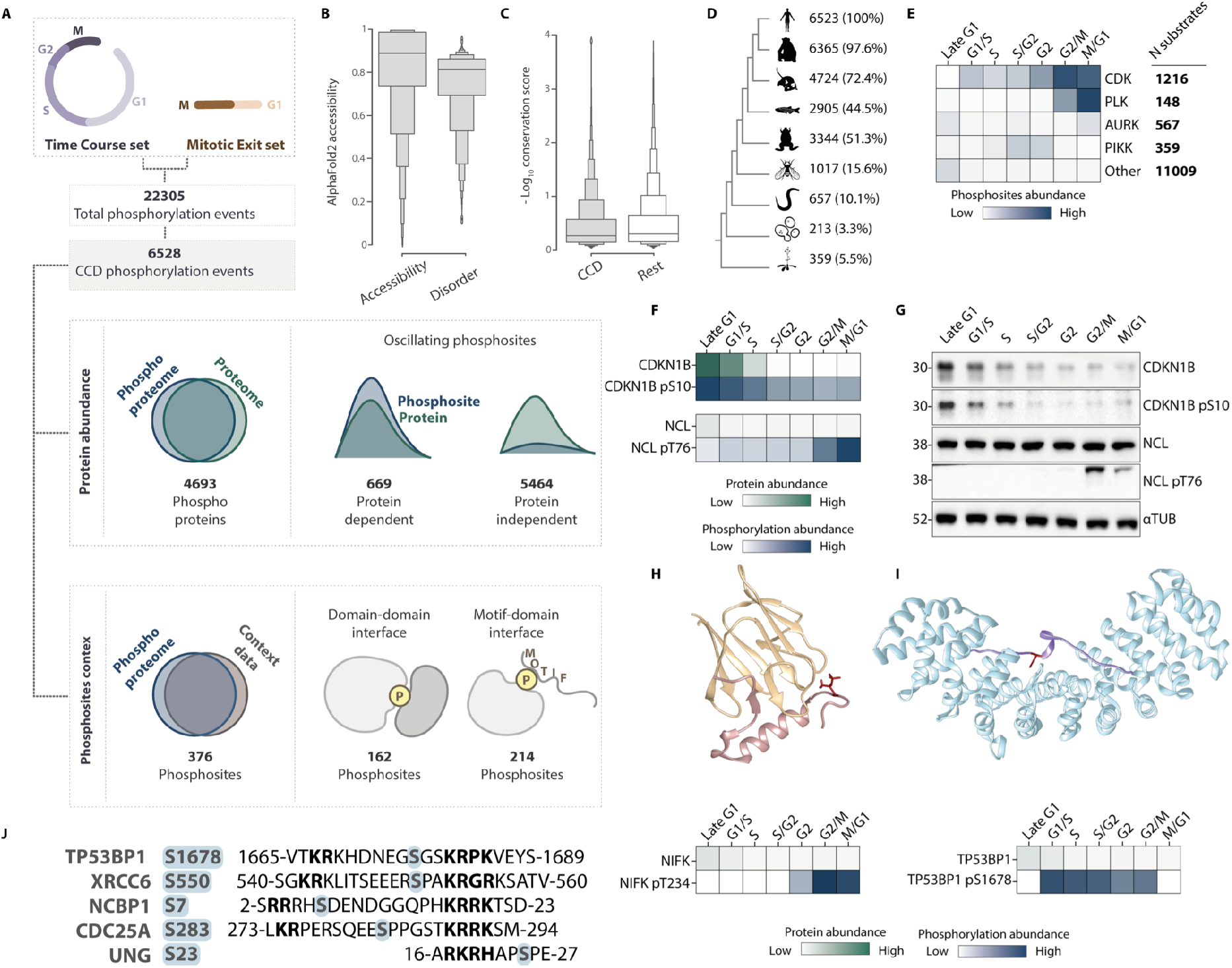
Cell cycle-dependent phosphorylation dynamics. **(A)** Workflow used to investigate phosphorylation dynamics during the cell cycle. CCD phosphorylation events detected in the Time Course and Mitotic Exit datasets were classified as protein-dependent and protein-independent based on their correlation with protein abundance. Furthermore, CCD phosphorylation events were annotated with structural features to define phosphorylation sites located at protein interfaces and phosphorylated residues that overlapped with known motifs. **(B)** Boxen plot comparing accessibility and disorder scores for CCD phosphorylation events. **(C)** Boxen plot showing differences in conservation scores between CCD and non-CCD phosphorylation events. **(D)** CCD phosphorylation sites conservation across different species. The numbers on the right represent the phosphorylation events identified in each species, and the percentage of enrichment is reported. **(E)** Heatmap showing the oscillation of phosphorylation sites predicted to be CDK, Aurora, PLK, PIKK and other kinases substrates. Phosphorylation changes are coloured according to their average abundance (log_2_ mean normalised values). Phosphorylation consensus motifs were obtained from Alexander *et al.,* 2011. **(F)** Heatmaps showing the log_2_ mean normalised protein and phosphorylation abundance changes during the cell cycle detected by MS. CDKN1B and NCL are shown as representative examples of protein-dependent and protein-independent phosphorylation events. **(G)** Western blot validation of protein abundance changes shown in (F). **(H)** Structure of MKI67 bound to NIFK (PDB ID: 2AFF). MKI67 is shown in yellow and NIFK in pink. This interaction is mediated by the phosphorylation of NIFK on Thr234, shown in red sticks. NIFK protein (green) and pThr234 (blue) abundance changes during cell cycle progression detected by MS are shown in the heatmap (bottom). Protein and phosphorylation changes are coloured according to their abundance (log_2_ mean normalised values). **(I)** Structure of KPNA2 bound to TP53BP1 (PDB ID: 6IU7). KPNA2 is shown in light blue and TP53BP1 in purple. This interaction is mediated by the phosphorylation of TP53BP1 on Ser1678, shown in red sticks. TP53BP1 protein (green) and pSer1678 (blue) changes during cell cycle progression detected by MS are shown in the heatmap (bottom). Protein and phosphorylation changes are coloured according to their abundance (log_2_ mean normalised values). **(J)** Examples of proteins containing motif consensus of phosphorylation sites adjacent to an NLS found in our dataset.

Phosphorylation dynamics during the cell cycle can be classified as protein-dependent and protein-independent, based on whether the phosphorylation patterns correlate with changes in protein abundance. To investigate these dynamics, we compared the oscillation profiles of the proteome and phosphoproteome using the CCD dataset (Figure 5A). We found that 669 phosphorylation sites exhibited protein-dependent oscillation, meaning that their phosphorylation dynamics mirrored protein changes throughout the cell cycle. For instance, we observed a rise in the phosphorylation of Ser10 on CDKN1B from late G1 to S phase closely mirroring the protein level changes (Figure 5F and 5G). This phosphorylation has been shown to regulate CDKN1B export from the nucleus to the cytoplasm for proteasomal degradation ^35,36^. In contrast, we identified 5,464 phosphorylation sites that exhibited protein-independent oscillation, not directly associated with changes in protein abundance. Proteins in this category show a higher degree of regulation at the phosphorylation level. For example, the protein abundance of the Origin Recognition Complex subunit 2 (ORC2) was almost stable throughout the cell cycle, but the levels of phospho-Thr266 increased from S to M phase (Supplementary Figure 6C). This phosphorylation event is known to be mediated by cyclin-dependent kinases to control the binding of ORC to chromatin and replication origins ^37^. Similarly, we identified nucleolin (NCL) to be stable throughout the cell cycle, however, several phosphorylation sites were significantly increasing in mitosis. We confirmed these findings by western blot showing the phosphorylation of Thr76 to increase rapidly in G2/M while the protein remained stable (Figure 5F and 5G).

To identify phosphorylation events involved in cell cycle regulation, we annotated the CCD phosphorylation sites located at protein interfaces and overlapping with known motifs (Supplementary Table 12). Among the 6,528 CCD phosphorylation events observed, we found 162 phosphorylation sites overlapping protein domain-domain interfaces and 214 phosphorylation sites overlapping with motifs (Figure 5A). One example of CCD phosphorylation sites specifically localised at protein interfaces is the Thr234 of NIFK known to interact with the FHA domain of Ki67 during mitosis ^38^. NIFK is initially phosphorylated by the nuclear kinase CDK1 on Thr238 and subsequently by the cytosolic kinase GSK3 on Thr234 ^39^. Importantly, we observed these phosphorylation sites to rise between the S and M phases of the cell cycle (Figure 5H), suggesting that the interaction between MKI67 and NIFK is time-regulated through changes in phosphorylation. An example of CCD phosphorylation sites not previously shown to oscillate during the cell cycle is the interaction between Importin subunit alpha-1 (KPNA2) and TP53-binding protein 1 (TP53BP1). This interaction is essential for the nuclear import of TP53BP1. Previous studies have shown that the phosphorylation of Ser1678 in TP53BP1, which is adjacent to a Nuclear Localisation Signal (NLS), is mediated by CDK1/cyclin B^40^. Furthermore, the substitution of Ser1678 with aspartate has been shown to reduce the binding affinity between TP53BP1 and KPNA2, leading to impaired nuclear import of TP53BP1 ^41^. We observed this phosphorylation site to peak from G1/S to G2/M phase, in agreement with the known role of TP53BP1 in DNA double-strand break repair during cell cycle progression (Figure 5I). Interestingly, CCD phosphorylation sites were observed flanking validated NLSes in several additional proteins involved in various cell cycle mechanisms (Figure 5J). For instance, we found that the X-ray repair cross-complementing protein 6 (XRCC6) exhibited phosphorylation on the Ser550, a site mitosis specific site matching a PLK consensus. This suggests that the oscillation of these phosphorylation sites may play a regulatory role in protein translocation during the cell cycle.

### Degradation mechanisms of CCD proteins

Ubiquitin-dependent protein degradation plays a key role in cell cycle progression ^42^. To investigate protein degradation mechanisms of CCD proteins, we cross-referenced the CCD proteomics dataset with a curated set of characterised degrons, a class of motif that recruits an E3 Ubiquitin ligase resulting in substrate ubiquitination and subsequent degradation. We found that 55 out of the 401 CCD proteins contained a previously described degron (Supplementary Table 13). Of these, 42 proteins contained experimentally validated APC/C recognition motifs, mainly D box and KEN box degrons. In most cases, these proteins displayed the canonical abundance profiles observed in APC/C substrates defined by a peak in abundance during mitosis, followed by a pronounced decrease by G1 ^43^ (Figure 6A). Within this protein subset, we identified mitotic cyclins (CCNB1, CCNA2) together with several key mitotic regulators (PTTG1/Securin) that have to be rapidly degraded to promote mitotic exit. Noteworthy, we also detected a subset of proteins containing experimentally validated degradation motifs, yet exhibiting non-canonical abundance pattern changes for APC/C substrates. These included proteins degraded before mitosis, such as FOXM1, CDC6 and CDC25A, previously shown to have a dual mechanism of degradation requiring the activity of both the APC/C and SCF to finely modulate their expression during the cell cycle ^44–46^. Furthermore, several validated APC/C degron-containing proteins remained stable during mitosis, without showing a decrease in abundance during G1 phase (CDC25C, DLGAP5) or were not found to oscillate in our dataset (GLS, ID1). We observed distinct behaviours within several closely related proteins. For instance, while PLK1 exhibited high oscillation, PLK2 remained stable throughout the cell cycle. We speculate this difference can be attributed to the absence of a degradation motif in PLK2 (Supplementary Figure 7A). Among the 55 proteins identified in the CCD dataset with known degradation mechanisms, 13 contained degradation motifs that were not recognised by the APC/C and displayed a distinct oscillation pattern compared to APC/C substrates. These include proteins with PIP degron (CDKN1A, KMT5), β-TrCP degron (WEE1, UHRF1, BORA) and Fbw7 degron (CCNE2, UNG) motifs. Interestingly, for the phosphorylation-dependent Fbw7 degron in Uracil-DNA Glycosylase (UNG), we detected the phosphorylation on Thr60 and Ser64 (Supplementary Figure 7B) previously shown to promote its degradation suggesting a delay between phosphorylation and degradation ^47^.

**Figure 6:**
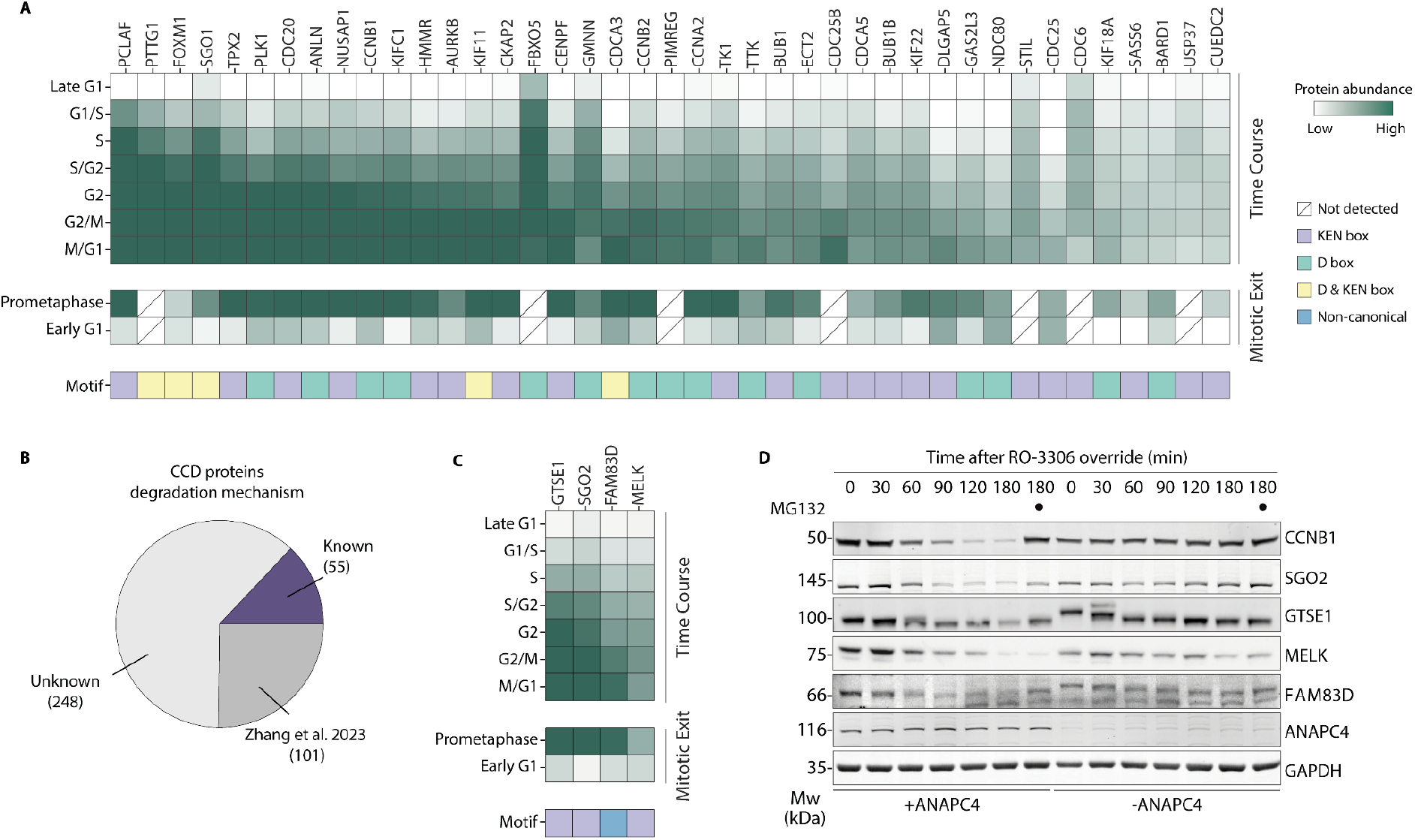
Investigating CCD protein degradation. **(A)** Heatmap showing canonical protein abundance profile for experimentally validated APC/C recognition motifs in both Time Course and Mitotic Exit datasets. Protein changes are coloured according to their abundance (log_2_ palbociclib arrest normalised values). Degradation motifs are indicated with different colours. Proteins not detected are shown with a dashed box. **(B)** The proportion of CCD proteins with experimentally validated APC/C recognition motifs and established degradation mechanisms (purple) versus those without a known mechanism (light grey). Proteins found to contain unstable peptides in a global peptide stability dataset ^48^ are also shown in dark grey. **(C)** Heatmap showing protein abundance changes for CCD proteins predicted to contain a degradation motif with high confidence and selected for validations. Protein changes are coloured according to their abundance (log_2_ palbociclib arrest normalised values). **(D)** Representative western blot analysis (n=3) of predicted mitotic APC/C substrates degradation in Hela Kyoto cells during the override of a mitotic checkpoint arrest by the CDK1 inhibitor RO-3306 in the presence and absence of ANAPC4. SGO2, GTSE1, MELK, FAM38D and known APC/C substrates (CCNB1) are stabilised by proteasome inhibition with MG132.

Out of the 401 CCD proteins identified in this study, only a small subset of 55 were found to contain experimentally validated degradation motifs (Figure 6B). As a result, the mechanism of degradation for the remaining 346 proteins is still unknown. By integrating proteome-wide peptide stability data into our dataset ^48^, we found that 101 of the proteins without a validated degron also contain peptides that are unstable in global peptide stability assays ^48^, suggesting they are likely to include degradation signals (Supplementary Table 13). The 346 proteins that do not possess known degradation motifs were screened using the PSSMSearch tool ^49^ to predict novel degradation motifs. This bioinformatic approach uncovered 60 regions with high similarity to the consensus of degron motifs known to modulate protein abundance in a cell cycle-dependent manner (Fbw7, β-TrCP, D box, KEN box, ABBA, PIP degrons) (Supplementary Table 13). To validate our predictions, we selected four CCD protein candidates (Figure 6C) based on statistically robust predictions and other high-confidence attributes such as degron accessibility and match to the canonical motif consensus. Since we previously showed that the depletion of ANAPC4, a crucial subunit of the APC/C results in the stabilisation of known APC/C substrates ^50^, we used this approach to validate novel candidates assessing protein stability upon mitotic release in ANAPC4-depleted or unperturbed HeLa cell extracts by western blot analysis (Figure 6D, Supplementary Figure 7D). Among the candidates showing strong motif predictions and protein abundance profiles consistent with the canonical APC/C substrate pattern, we confirmed that SGO2 is targeted for degradation by the APC/C during early mitosis and follows a degradation pattern similar to the well-known substrate CCNB1, in agreement with our previous findings ^50^. However, further experiments are required to validate the SGO2 degradation motif, which we predicted to be a KEN box. Next, we found that GTSE1 accumulated upon APC4 depletion or proteasome inhibition, confirming that it is a mitotic APC/C substrate. Of note, mouse GTSE1 has been previously identified as a KEN box-dependent APC/C substrate ^51^, indicating that this degradation motif is evolutionarily conserved. Within this group we also found proteins, such as MELK, that exhibited partial stabilisation upon ANAPC4 depletion, suggesting that their degradation may be regulated by multiple degradation pathways. Additionally, we identified protein candidates with strong motif predictions that did not follow the canonical APC/C substrate abundance pattern. For example, CHAF1B increased in abundance through the cell cycle but did not exhibit a significant decrease upon prometaphase release in the Mitotic Exit dataset. We found that the degree of depletion we achieved for ANAPC4 was not sufficient to observe stabilisation of CHAF1B, suggesting that it is not regulated through the APC/C but rather through alternative degradation mechanisms that are yet to be defined (Supplementary Figure 7C). Finally, the majority of our predicted candidates showed protein CCD abundance profiles consistent with APC/C substrates, yet, they did not contain an obvious APC/C degron. For instance, the spindle-associated protein FAM83D ^52^ exhibited a significant accumulation in ANAPC4-depleted cells (Figure 6D) and a protein oscillation consistent with the APC/C substrate pattern (Supplementary Figures 7E and 7F). This observation aligns with other examples of APC/C substrates that lack canonical degradation motifs, such as AURKA and SGO1 which contain a non-canonical D box ^53,54^. This set of proteins with non-canonical degradation motifs raises intriguing questions about the intricacies underlying their degradation mechanisms.

### Database

The data produced and collected in this study is available in the Cell Cycle Database (CCdb) resource (see Supplementary Document S2 for a detailed description). The database holds the cell cycle abundance changes for proteins and phosphorylation sites for the Time Course, Mitotic Exit and Serum Starvation datasets, the respective statistical metrics and the cell cycle dependency status. The data is additionally enriched with external information on protein, phosphorylation site, mRNA abundance and cell cycle dependency from publicly available datasets. Protein level information on stability and degrons is also provided. For phosphorylation sites, annotations on the accessibility, domain and motif overlap, and presence in other publicly available phosphoproteomic datasets are annotated. The CCdb links cell cycle-dependent abundance dynamics to functional changes illuminating the biological role of oscillating proteins and phosphorylation events and is available at https://slim.icr.ac.uk/cell_cycle/.

## Discussion

In this study, we present a comprehensive analysis of protein and phosphorylation changes during the human cell cycle in non-transformed cells. We have characterised a set of 401 cell cycle oscillating proteins with high-temporal resolution and coverage using deep quantitative mass spectrometry. This cell cycle signature set includes 190 well-known proteins, along with 211 proteins newly found to oscillate during the cell cycle, thus providing a valuable resource for researchers in this field (Supplementary Table 8). Furthermore, we developed the Cell Cycle database (CCdb; https://slim.icr.ac.uk/cell_cycle), a user-friendly web-based resource that allows for interactive exploration of our comprehensive dataset, thereby providing a valuable tool for the cell cycle research community.

We aimed to address the limitations of previous research by providing a comprehensive understanding of the protein and phosphorylation changes that occur throughout the cell cycle. Existing datasets have primarily focused on cancer cell lines (Hela ^7–9^, NB4 ^5^, T98G ^6^), which may not accurately reflect the normal cell cycle progression ^55,56^. Instead, we used RPE-1 cells that contain a normal repertoire of cell cycle regulators and carefully optimised the conditions to induce robust synchrony while minimising the impact on the cell cycle ^14^. Previous studies have used chemical synchronisation methods such as double thymidine block known to cause DNA damage and chromosomal rearrangements ^10–12^ or serum starvation that has been shown to induce cell stress and semi-synchronous re-entry into cell cycle progression ^57,58^. Moreover, FACS-based methods have drawbacks in terms of protein coverage and temporal resolution due to the limited number of cells obtained and the parameters used to collect different cell cycle stages. To overcome these limitations, we used palbociclib-induced synchronisation to investigate global changes throughout the cell cycle. Palbociclib is a CDK4/6 inhibitor shown to efficiently arrest cells before the natural restriction point and generate highly synchronised cell cycle progression upon release ^12,13^. Although it is important to acknowledge palbociclib synchronisation may influence cell cycle timing and impact the protein and phosphorylation profiles observed in this study, we found that using palbociclib arrest is an effective approach to collect a large population of cells in specific phases of the cell cycle, in line with previous reports ^12,13^. However, we observed reduced synchrony due to their natural progression through different cell cycle phases towards the end of the cell cycle. Therefore, we generated an additional dataset specifically focusing on events occurring at mitotic exit by inducing prometaphase arrest with the Eg5 inhibitor DMA and subsequent release into early G1.

Previous studies have identified several cell cycle regulators using MS-based proteomics. However, when comparing existing datasets only a small number of key cell cycle regulators, such as CCNB1, were consistently detected across different studies (Supplementary Figure 1D). Moreover, due to technical advancements in MS over the last decades, cell cycle proteomics and phosphoproteomics datasets have improved in resolution and allowed for a more precise representation of cell cycle dynamics. As the most recent study, our dataset provides an unprecedented resolution, covering the major cyclins as well as key regulatory proteins throughout critical phases of the cell cycle. We define a set of regulatory proteins with oscillating abundance patterns throughout the cell cycle as cell cycle-dependent (CCD) proteins (Supplementary Table 8). These include well-known and novel proteins whose cell cycle dynamics were not previously reported or clearly understood. One example is the oscillation of CCND3, showing opposite patterns to CCND1 (Figures 2D and 2E). Another intriguing finding is the oscillation of CDKs, which rises from the S to G2/M phase (Figure 4D, left panel) and contradicts the conventional knowledge that CDKs remain constant throughout the cell cycle but is consistent with previous research showing that CDK1 is regulated at the transcription level ^15,25^. Overall, the majority of the detected proteins remained stable throughout the cell cycle, highlighting that protein abundance represents only one layer of the complex regulation.

Our comprehensive dataset provides a valuable opportunity to study the regulation of protein degradation during the cell cycle. We not only identified 52 proteins with a well-established cell cycle-dependent degradation mechanism but also revealed several novel potential substrates of the anaphase-promoting complex/cyclosome (APC/C), a key regulator of protein degradation during mitosis (Supplementary Table 13). We investigate the degradation mechanisms for a set of proteins that we predicted to contain uncharacterised degradation motifs. Among these, we confirmed that GTSE1, MELK, and FAM83D are targeted for degradation by the APC/C during mitotic exit (Figure 6C and 6D). Interestingly, the majority of the CCD proteins do not have any clear validated or predicted degradation mechanism, suggesting they are regulated through novel modes of protein degradation. One possibility is that there may be variations of the canonical degradation motifs not yet identified, as most cell cycle-associated degradation motifs (Fbw7, β-TrCP, D box, KEN box, ABBA, PIP degrons) remain poorly characterised. Another explanation is that these proteins may contain a novel class of degradation motif. Alternatively, protein degradation might occur through interactions with scaffolded complexes through other proteins containing a degradation motif.

While substantial progress has been made in mapping the cell cycle proteome, there is still a considerable lack of knowledge regarding the complexity of the cell cycle phosphoproteome ^8^. (Figure 2M, Supplementary Figure 3H). Despite technological advancements in the identification of phosphorylations using MS, many cell cycle phosphorylation events remain uncovered due to their transient nature and technical limitations. This is evident in the limited overlap of the existing cell cycle phosphoproteomic datasets and highlights that the CCD phosphoproteome is far from being fully elucidated. Our study uncovers 6,528 CCD phosphorylation events, 5,464 of which oscillate independently of the protein oscillation underlining the critical role played by phosphorylation in regulating protein function throughout the cell cycle. Of note, the vast majority of these phosphorylation events had not previously been reported to oscillate during cell cycle progression (Figure 2M). These phosphorylation events include several well-established regulatory phosphorylations, such as the inhibitory and activating phosphorylation of CDK1 (Figure 2H and 2I), and numerous CCD phosphorylation sites overlapping known protein interaction interfaces (figure 5A). Overall, the functional role of most CCD phosphorylation events is currently uncharacterised and poses an intriguing opportunity for further investigation considering the potential cell cycle stage-dependent regulation encoded by these sites.

To investigate the interplay between cell cycle regulation and other biological processes, we incorporated data from external studies. Combining protein oscillation with data on protein localisation and protein complexes (Supplementary Table 10), we observed differential stability that explains cell cycle biology. For example, CDC20, a regulatory protein crucial for the activation of the APC/C, was found to be highly oscillating compared to other components of the complex, in line with its importance in controlling the timing of the cell cycle by targeting specific substrates for degradation and exit from mitosis (Figure 4C). Moreover, the integration of the cell cycle-regulated transcriptome ^15^ (Supplementary Table 11) with our proteomics data revealed that roughly half of the CCD proteins exhibit oscillating transcripts, whilst the other half maintain stable transcripts (Figure 4E). This observation suggests that there is a complex regulatory network governing the oscillation of CCD proteins during the cell cycle on both the transcriptional, post-transcriptional and post-translational levels. Furthermore, by integrating our CCD proteomics dataset with gene essentiality data ^28^, we identified several key cell cycle regulators that are essential for cell viability. In contrast, a minor proportion of CCD proteins were not essential, raising questions about their functional significance in the cell cycle and the possibility of redundancy or compensatory mechanisms.

In summary, we present a comprehensive proteomic and phosphoproteomic analysis of the human cell cycle in non-transformed hTERT-RPE-1 cells with high-temporal resolution and coverage. We provide a detailed view of the dynamic changes in the human proteome and phosphoproteome throughout the cell cycle. These data have been made accessible in the Cell Cycle database (CCdb; http://slim.icr.ac.uk/cell_cycle), a web-based resource for the cell cycle community. This cell cycle dataset, along with the associated exploration tools, holds significant potential for diverse applications and provides valuable insights for researchers exploring this fundamental cellular process.

## Methods

### Cell lines

Human Retinal Pigment Epithelial hTERT (RPE-1) FRT and hTERT RPE-1 mRuby-PCNA cells were grown in DMEM/F12 medium supplemented with 10% (v/v) foetal bovine serum (FBS), 1% (v/v) Glutamax, 0.348% sodium bicarbonate and 1% (v/v) penicillin-streptomycin. HeLa Kyoto and HeLa FRT/TR 3xFlag-Venus-SBP-APC4 + NLS-TIR1/NES-TIR1 + mAID-nanobody cells were grown in DMEM supplemented with 10% (v/v) FBS, 1% (v/v) penicillin-streptomycin, 1% (v/v) Glutamax, 0.5 mg/ml Amphotericin B.

### Cell synchronisation

For the Time Course dataset, cells were seeded at a confluency of 4.4×10^3^/cm^2^ overnight. Then, palbociclib was added at a final concentration of 150 nM. Upon 24h incubation with the inhibitor, cells were washed three times with phosphate-buffered saline (PBS) buffer, released in fresh medium and subsequently collected at different time points: 0h (Late G1 palbociclib arrest), 2h (late G1 phase), 4h (G1/S phase), 6h (S phase), 8h (S/G2 phase), 10h (G2 phase), 12h (G2/M phase) and 15h (M/early G1 phase). For the Mitotic Exit dataset, cells were first released from palbociclib synchronisation for 10h (G2 phase) and then incubated with Dimethylenastron (DMA) at 1μM final concentration. After 4h, mitotic cells were harvested by shake-off and washed four times in PBS. Then, cells were released in fresh medium and harvested at two time points: 0h (prometaphase arrest) and 5h (early G1 phase). For the Serum Starvation dataset, 2.5 M cells were seeded in 150mm plates overnight. The next day, cells were washed three times with PBS and incubated with serum free medium for 24h. Cells were then released in complete medium and harvested at two time points: 0h (G0 phase) and 8h (G1 phase). Synchronised cells were harvested at the indicated time points, washed twice with PBS and stored at - 80°C before mass spectrometry analysis. Aliquots from each time point were fixed for flow cytometry analysis.

For APC/C substrates validations, HeLa Kyoto cells were synchronised in early S phase by a double thymidine block and release protocol. Briefly, cells at 40% confluency were treated with 2.5 mM Thymidine for 19h, followed by a release into fresh medium for 12h and a second thymidine block for 16h. After the second release cells were collected at the indicated time points.

### Sample preparation for proteomics and phosphoproteomics analysis

Pellets from synchronised cells were lysed in 150 μL lysis buffer of 100 mM triethylammonium bicarbonate (TEAB), 1% sodium deoxycholate (SDC), 10% isopropanol, 50 mM NaCl and Halt protease and phosphatase inhibitor cocktail on ice, with 15 sec of pulsed probe sonication followed by heating at 90°C for 5 min and another round of sonication for 5 sec. Protein concentration was measured with the Quick Start Bradford protein assay according to manufacturer’s instructions. Protein aliquots of 60 μg were reduced with 5 mM tris-2-carboxyethyl phosphine (TCEP) for 1 h at 60 °C and alkylated with 10 mM iodoacetamide (IAA) for 30 min in the dark. Proteins were digested overnight with trypsin at 75 ng/μL final concentration. Peptides were labelled with the TMTpro-16plex reagents (Thermo Fisher Scientific) according to manufacturer’s instructions. The pooled sample was acidified with 1% formic acid (FA), the precipitated SDC was removed by centrifugation and the supernatant was SpeedVac dried.

### High-pH Reversed-Phase peptide fractionation

Peptides were fractionated with high-pH Reversed-Phase (RP) chromatography using the XBridge C18 column (2.1 x 150 mm, 3.5 μm, Waters) on a Dionex UltiMate 3000 HPLC system. Mobile phase A was 0.1% (v/v) ammonium hydroxide and mobile phase B was acetonitrile, 0.1% (v/v) ammonium hydroxide. The TMTpro labelled peptides were fractionated at 0.2 mL/min with the following gradient: 5 min at 5% B, up to 12% B in 3 min, linear gradient to 35% B in 32 min, gradient to 80% B in 5 min, isocratic for 5 min and re-equilibration to 5% B. Fractions were collected every 42 sec, combined in 30 fractions and SpeedVac dried.

### Phosphopeptide enrichment

Phosphopeptide enrichment was performed in the first 24 peptide fractions with the High-Select Fe-NTA Phosphopeptide Enrichment Kit using a modified protocol in a well plate tip-array format. A volume of 50 μL resin/buffer was transferred on top of 10 μL filter tips that were fitted on a 96-well plate using a tip rack. The resin was washed three times with 40 μL wash/binding solution and centrifugation at 500 g for 1 min. Peptides were reconstituted in 30 μL wash/binding solution and loaded onto the tip-columns with the resin. After 30 min, the resin was washed three times with wash/binding solution and the flow-throughs were collected in a clean 96-well plate with centrifugation at 500 g for 1 min each time. Phosphopeptides were eluted twice with 40 μL elution buffer in a clean 96-well plate with centrifugation at 500 g for 1 min, transferred in glass vials and SpeedVac dried.

### Liquid Chromatography (LC) Mass Spectrometry (MS) analysis

LC-MS analysis was performed on a Dionex UltiMate 3000 UHPLC system coupled with the Orbitrap Lumos Mass Spectrometer (Thermo Fisher Scientific). Peptides were loaded onto the Acclaim PepMap C18 trapping column (100 μm × 2 cm, 5 μm, 100 Å) at flow rate 10 μL/min and analysed with an Acclaim PepMap (75 μm × 50 cm, 2 μm, 100 Å) C18 capillary column connected to a stainless-steel emitter. Mobile phase A was 0.1% FA and B was 80% acetonitrile, 0.1% FA. For the phosphopeptide analysis, the separation method was: 60 min linear gradient 5%-38% B at flow rate 300 nL/min. For the flow-through analysis a 90 min gradient 5%-38% B was used. MS scans were acquired in the range of 375-1,500 m/z with mass resolution of 120k, AGC 4×10^5^ and max IT 50 ms. Precursors were selected with the top speed mode in 3 sec cycles and isolated for HCD fragmentation with quadrupole isolation width 0.7 Th. Collision energy was 36% with AGC 1×10^5^ and max IT 100 ms (or 86 ms for flow-throughs) at 50k resolution. Targeted precursors were dynamically excluded for further fragmentation for 30 seconds (or 45 sec for flow-throughs) with 7 ppm mass tolerance.

### Flow cytometry analysis

Cells were fixed by adding ice-cold 70% ethanol in PBS dropwise to the pellet while vortexing and stored at −20°C prior to staining. Fixed cells were washed twice with PBS and incubated with propidium iodide and RNase at 37°C for 20 min in the dark. Stained cells were analysed on a BD LSR II flow cytometer and data acquired using BD FACSDiva software. Data was analysed using Flowjo software.

### ANAPC4 depletion assay

To deplete endogenous ANAPC4, HeLa FRT/TR 3xFlag-Venus-SBP-ANAPC4 + NLS-TIR1/NES-TIR1 + mAID-nanobody cells were synchronised at the beginning of S phase with a 2.5 mM thymidine for 24h and released into fresh medium containing 0.5 mM synthetic auxin analog 1-Naphthaleneacetic acid (NAA) and 245 nM taxol for a total of 13 h. Then, NAA-treated (-ANAPC4) and non-treated (+ANAPC4) prometaphase cells were collected by mitotic shake-off, washed, re-suspended in fresh media containing 9 μM RO-3306 and 10 μM proteasome inhibitor MG-132 as indicated to induce mitotic exit. After the indicated time, cells were harvested, washed twice in PBS, snap frozen in liquid nitrogen and stored at −80°C before further analysis.

### Western blot analysis

For total protein and phosphorylation changes validations, cell pellets were lysed in extraction buffer (1% (v/v) Triton X-100, 10 mM Tris-HCl, pH 7.4, 5 mM EDTA, 50 mM NaCl, 50 mM sodium fluoride, 2 mM Na3VO4, supplemented with cOmplete™ EDTA-free Protease Inhibitor Cocktail and Phosstop™-phosphatase inhibitor tablets) by passage through a 26-gauge needle six times. The lysate was incubated on ice for 5 min and then clarified by centrifugation (14,000 rpm for 10 min at 4 °C). For ANAPC4 depletion assay, cell pellets were lysed in extraction buffer (PBS/1% Triton-X100 supplemented with cOmplete™ EDTA-free Protease Inhibitor Cocktail, Phosstop™-phosphatase inhibitor tablets and 1 mM DL-Dithiothreitol (DTT)). The lysate was incubated on ice for 25 min, sonicated for 20s to shear DNA, boiled for 10 min at 95°C and then clarified by centrifugation (21,000 rcf for 10 min at 4°C). The protein concentration was quantified with a Bradford assay. Proteins were resolved by SDS-PAGE and transferred onto a nitrocellulose or PVDF membrane for immunoblot analysis. After 1 h blocking with 5% milk in TBST at room temperature (RT) for 1 h, membranes were incubated overnight at 4°C with the following primary antibodies: CCND1 (Santa Cruz Biotechnology, sc-2004), CCND3 (Santa Cruz Biotechnology, sc-6283), CCNE2 (Santa Cruz Biotechnology, sc-248), CCNA (Santa Cruz Biotechnology, sc-271682), CCNA (Santa Cruz Biotechnology, sc-596), CCNB1 (Santa Cruz Biotechnology, sc-245), GAPDH (Santa Cruz Biotechnology, sc-51907), CDK1 pT161 (Cell Signalling Technology, 9114), CDK1 pY15 (Cell Signalling Technology, 9111), Alpha Tubulin (Abcam, ab52866), PLK1(Millipore, 05-844), CDK2 (Santa Cruz Biotechnology, sc-163), CDK1 (Abcam, ab32094), CDC20 (Santa Cruz Biotechnology, sc-13162), UBR7 (Thermo Fisher Scientific, A304-130A), UBE2S (Cell Signalling Technology, 11878), CDKN1B (Dako, M7203), CDKN1B pS10 (Santa Cruz Biotechnology, sc-12939), NCL (Abcam, ab22758), NCL pT76 (Abcam, ab168363), CCNB1 (Santa Cruz Biotechnology, sc-245), SGO2 (Thermo Fisher Scientific, A301-261A), GTSE1 (Abnova, H00051512-B01P), MELK (Thermo Fisher Scientific, A303-136A), FAM83D (Custom antibody from Erich Nigg), ANAPC4 (Thermo Fisher Scientific, A301-175A), Phospho-CDK substrate motif [(K/H)pSP] (Cell Signalling Technology, 9477), JUNB (Proteintech, 10486-1-AP), CHAF1B (Thermo Fisher Scientific, A301-085A), Histone H3 pS10 (Cell Signalling Technology, 3377), Actin (Sigma-Aldrich, A5441). After washed with TBST for three times, membranes were incubated for 1h at RT with the following secondary antibodies: IRDye 680RD Donkey anti-Rabbit IgG (H+L) (LI-COR Biosciences, 926-68073), IRDye 800CW Donkey anti-Rabbit IgG (H + L) (LI-COR Biosciences, 926-32213), IRDye 800CW Donkey anti-Mouse IgG (H + L) (LI-COR Biosciences, 926-32212), IRDye 680RD Donkey anti-Rabbit IgG (H + L) (LI-COR Biosciences, 926-68073), anti-mouse HRP (Cell Signalling Technology, 7076), anti-rabbit HRP (Cell Signalling Technology, 7074). After washing three times with TBST, blots were detected with an Odyssey infrared scanner or SuperSignal™ West Pico PLUS Chemiluminescent Substrate on an ChemiDoc™ MP Imaging System.

### Immunofluorescence

hTERT RPE-1 mRuby-PCNA cells were seeded on glass cover glasses. After 24h, cells were fixed for 5 min in 4% paraformaldehyde in PBS at RT, permeabilised in 0.1% Triton X-100, 0.02% SDS in PBS for 5 min at RT, blocked in blocking buffer (2% albumin fraction V (Roth, 8076.4) in PBST) for 1h at RT, and incubated for 1h with the following primary antibodies: CCNB1 (Santa Cruz Biotechnology, sc-245), FAM83D (Custom antibody from Erich Nigg). Afterwards, cover glasses were washed 3 times in PBST, incubated in blocking buffer supplemented with 1 µg/ml Hoechst 33342 for 1h at RT with secondary antibodies: Alexa Fluor 647 goat anti-mouse (Invitrogen, A-21236), Alexa Fluor 488 chicken anti-rabbit (Invitrogen, A-21441). Then, washed 3 times in PBST, post-fixed in 4% paraformaldehyde, washed once in PBST, and mounted on glass slides using Vectashield mounting medium. Digital images were captured on a Delta Vision wide-field deconvolution fluorescence microscope equipped with a PCO Edge/sCMOS camera (Imsol) at binning=2, and a Olympus Plan Apo N 60X/NA 1.42 lens (Olympus) objective.

### MS data processing

The mass spectra were processed using Proteome Discoverer 2.4 (Thermo Scientific) with the Sequest HT search engine for peptide identification and quantification. The precursor and fragment ion mass tolerances were 20 ppm and 0.02 Da respectively. Spectra were searched for fully tryptic peptides with a maximum of 2 missed-cleavages. TMTpro at N-terminus/K and Carbamidomethyl at C were set as static modifications. Oxidation of M, Deamidation of N/Q and Phosphorylation of S/T/Y were set as dynamic modifications. Spectra were searched against reviewed UniProt Homo sapiens protein entries (version 20apr21), peptide confidence was estimated with the Percolator node and peptides were filtered at q-value < 0.01 based on target-decoy database search. The consensus search result was filtered to a protein false discovery rate adjusted (FDR) of 0.01 (High) and 0.05 (Medium). External contaminants were removed from protein lists before further analysis. The reporter ion quantifier node included a TMTpro quantification method with an integration window tolerance of 15 ppm. Only peptides with average reporter signal-to-noise > 3 were used, and phosphorylation localisation probabilities were estimated with the IMP-ptmRS node. Abundances from Proteome Discoverer were processed in Python and R for the Time Course, Mitotic Exit and Serum Starvation datasets. The protein and phosphorylation abundances were normalised by dividing the median intensity for each sample to account for changes in the total cell content during the cell cycle. Abundances of phospho-peptides that corresponded to the same phosphorylation site in the same protein were added to produce a single abundance for each phosphorylation site. Protein and phosphorylation site abundances were subsequently normalised using further approaches to allow specific analyses (described in Supplementary Document 1).

### Defining cell cycle-dependent (CCD) proteins and phosphorylation sites

To define significant protein and phosphorylation changes in the Time Course dataset, we performed one-way ANOVA analysis and corrected p-values using the Benjamini-Hochberg method for multiple hypothesis testing. We then used a polynomial curve to model the oscillation patterns of each protein and phosphorylation site abundance. The *0-max normalised* abundance was modelled by a polynomial of degree 2 curve, where *y* encoded the abundance and x encoded the time points in incrementing steps size 1. The curve was calculated using *numpy.poly1d* method from the *numpy* library and curve fold change was calculated by dividing the maximum value of the new curve fitted abundance by the minimum curve fitted abundance. Fitted abundances below zero were given a pseudo-abundance of 0.05. Cell cycle-dependent (CCD) proteins and phosphorylation sites were defined for the Time Course dataset based on GO term enrichment analysis on the term “cell cycle” (GO:0007049). Protein curve fold change cut-offs were defined by quantifying the enrichment of the “cell cycle” term at a range of curve fold change or fold change cut-offs in increments of 0.1. Each analysis compared the proteins above that cut-off against all remaining proteins observed in mass spectrometry data (Supplementary Table 3). Cut-offs were selected so that the CCD set had a significant enrichment of the “cell cycle” term with a p-value less than 0.01. For the Time Course dataset, a curve fold change cut-off of 1.2 (i.e. at least 20% difference between the minimum and maximum abundance) was determined and applied, and further filtering based on adjusted ANOVA p-value ≤ 0.01 and FDR confidence (FDR ≤ 0.05 for proteins and peptide FDR ≤ 0.01 for phosphorylation) were performed to define the CCD set.

To define significant protein and phosphorylation changes in the Mitotic Exit and Serum Starvation datasets, the three replicates were grouped based on their time points (DMA arrest, DMA release, serum starvation arrest, serum starvation release and palbociclib arrest). First, phosphopeptide abundances in the DMA Arrest arrest were correct to remove the bias introduced by the total intensity in this hyperphosphorylated time point as described in Supplementary Document 1. Next time points were paired and pairwise limma analyses were performed to generate fold change and adjusted p-values for each protein and phosphorylation site. Key pairs tested include the DMA arrest/release (Mitotic Exit) and Serum Starvation arrest/release. Protein fold change cut-offs were defined as described for the Time Course dataset. For the Mitotic Exit dataset, a log_2_ fold change cutoff of 0.5 was determined and applied for proteins and a cutoff of 1 was chosen for phosphorylation sites. The more conservative cut-off for phosphorylation sites reflects the large phosphorylation abundance changes observed in the Mitotic Exit set. Proteins and phosphorylation sites were further filtered with adjusted limma p-value ≤ 0.01 to define the CCD set.

Stable proteins and phosphorylation sites were defined based on the Time Course data as those with: (i) a standard deviation ≤ 0.05; (ii) an adjusted ANOVA p-value > 0.001; and an FDR ≤ 0.05 for proteins and peptide FDR ≤ 0.01 for phosphorylation sites. The standard deviation was calculated on the *0-max normalised* abundance data. Finally, for both the Time Course and Mitotic Exit sets, protein-dependent and protein-independent phosphorylation abundance changes were defined. CCD phosphorylation sites in CCD proteins were characterised as “CCD phosphorylation site - protein-dependent”, whereas CCD phosphorylation sites in non-CCD proteins were characterised as “CCD phosphorylation site - protein-independent.”.

### Cluster analysis for protein abundance profiles

Protein abundance profile clustering was performed with the T-distributed Stochastic Neighbour Embedding (TSNE) approach and using the *sklearn.manifold.TSNE* function from the Scikit-learn library. The TSNE approach converts high-dimensional data into lower dimensions, allowing relationships in complex datasets to be easily extracted. The *min-max* normalised abundance was calculated on the Time Course CCD protein set. The *min-max* abundance normalisation defines the minimum value as 0 and the maximum value as 1, and permits comparison of abundance profiles independent of the amplitude of the abundance change. The 16 *min-max* abundances were collapsed to one dimension. The distance of each protein in the Time Course CCD protein set was calculated to a set of 6 well-known cell cycle markers: cyclin D1, cyclin E2, MYB, Geminin, cyclin A2 and cyclin B1 and the protein was classified to the closest marker’s cluster.

### Protein annotation

A range of sources of data were used to functionally annotate the proteins quantified during the analysis. General protein information was retrieved from the UniProt resource. For protein localisation, “Cellular Component” GO term annotations were retrieved from the QuickGO Gene Ontology and Gene Ontology annotation database ^59^. Protein members of complexes were retrieved from the Complex Portal ^21^. Three key regulatory protein classes were defined as follows: (i) Proteins containing a Pfam kinase domain were defined as kinases ^60^; (ii) Proteins matching with the GO Term “DNA-binding transcription factor activity” (GO:0003700) were defined as transcription factors; and (iii) E3 ubiquitin ligases were derived from manual curation. Protein stability information on the mean protein half-life and the proportion of 0h protein remaining 8h after cycloheximide addition was retrieved from cycloheximide chase coupled to mass spectrometry experiments by Li *et al* ^27^. A set of validated degron motifs (degrons) was collected, this included APC/C degrons from Davey *et al* ^61^, non-APC/C degrons from the ELM database ^62^ and manually curated motifs. A set of peptides containing sequence-dependent degrons was collected from Zhang *et al* ^48^, peptides were annotated by Peptools and inaccessible peptides (AlphaFold Accessibility < 0.5) were discarded. The Mean Gene Effect scores from a CRISPR gene knockout and cell viability assays were retrieved from the Cancer Dependency Map resource ^28^. Data on a protein’s association with and role in cancer was retrieved from the Cancer Gene Census (CGC) database (https://cancer.sanger.ac.uk/census) and 299 DRIVE cancer driver genes derived from a PanCancer and PanSoftware analysis were retrieved for ^63^.

### Phosphorylation site annotation

The phosphorylation sites quantified during the analysis were annotated with a range of relevant data. Phosphorylation site accessibility and disorder scores are calculated from the DSSP residue accessible surface area (ASA) of AlphaFold2 models ^64^. Accessibility is the surface accessibility score derived from AlphaFold2 models for the single phosphorylated residue normalised by the maximum possible accessibility for that residue (described in Supplementary Document 1). The disorder score is a windowed version of the accessibility score, disordered regions contain many contiguous highly accessible residues, and is calculated as the mean accessibility for a region overlapping a given site with flanking regions of 15 amino acids. The taxonomic range of each phosphorylation site was calculated using the PepTools peptide annotation tool ^31^. The taxonomic range was derived from orthologue alignments created using the GOPHER algorithm against the Quest of Orthologs (QFO) protein set. Each phosphorylation site was annotated based on their conservation in the following model organisms H.sapiens, P.troglodytes, M.musculus, D.rerio, X.tropicalis, D.melanogaster, C.elegans, S.cerevisiae and A.thaliana. A phosphorylation site was defined as conserved in a species when a serine, threonine or tyrosine was present in the same position of the alignment as the human site. The phosphorylation site functional region overlap was tested on two datasets: (i) domain-domain interfaces and (ii) domain-motif interfaces. The set of interaction interfaces was derived from the structurally characterised complexes from the PDB database ^65^. Interface residues were determined as residues with heavy atoms within less than 6 Å distance from another protein chain in structures of protein complexes with 2 or more subunits. Binding partners defined as motifs using in-house software were added to the domain-motif interfaces and the remaining were added to the domain-domain interfaces set. The majority of the domain-motif interfaces set of validated short linear motifs were derived from ELM ^62^ and in-house curated motifs. Finally, phosphorylation sites were annotated for similarity to the specificity determinants of key cell cycle-related kinases. Optimal substrate phosphorylation motifs for each kinase were defined as follows: PLK: “[DNE].[ST][FGAVLIMW][GAVLIPFMW]”, CDK: “[ST]P.[RK]”, Aurora kinase: “R.[ST][GAVLIFMW]” and PIKK “[ST]Q” ^66^. Each phosphorylation site was compared to the regular expressions and matches to one or more consensus was annotated.

### Enrichment analysis annotation

Classical enrichment analysis was performed using the PANTHER enrichment tool ^67^ to identify enriched functional annotations. The analysis includes GO term (Biological Process, Molecular Function and Cellular Component) and Reactome pathway annotation. Both the protein and phosphorylation datasets analyses are performed on the protein level. All MS data was split into CCD and non-CCD proteins or phosphorylation and CCD enrichment was calculated using Fisher’s Exact test with the non-CCD set as the background.

### Degrons predictions

Peptides from the ELM DEG (i.e. degron) functional classes were retrieved programmatically from the ELM database ^62^. The degron classes were categorised as CC Degron Class based on their known functional role in the cell cycle. The peptides for each class were aligned with their ELM consensus and converted to PSSM with the PSI-BLAST IC PSSM scoring method. The PSSMs were scanned against the CCD proteins using the PSSMSearch motif discovery software ^49^ with a PSSM p-value cutoff of 0.001. High Confidence Degron predictions were defined as all uncharacterised instances that belong to a CC Degron Class, had a PSSM p-value < 0.00005, accessibility > 0.5, and disorder score > 0.5.

### Publicly available cell cycle datasets

Proteomics and phosphoproteomics data were parsed from the supplementary material of the following studies: Ly ^5^, Ginno ^6^, Herr ^7^, Olsen ^8^, McCloy et al ^17^. and Becher ^9^. Protein abundance data from immunofluorescence experiments was parsed from the Human Protein Atlas ^15^. Transcriptomics data were parsed from Herr ^7^, Giotti ^68^, Bar-Joseph ^69^, Grant ^70^, Whitfield ^71^, and Pena-Diaz ^72^. Cell cycle transcriptomics data linked to FUCCI fluorescent cell cycle reporter expression was retrieved from ^15^. The TPMs (transcripts per million) for each protein were grouped according to their FUCCI reporter pseudo times into three time points (G1, S and G2/M) and the median TPM value for each time point was taken as the representative TPM. The time point with the maximum median TPM was considered the peak. The CCD status of the proteins and transcripts were taken directly from ^15^.

### Data availability

The mass spectrometry data have been deposited in the ProteomeXchange Consortium ^73^ and are publicly available as of the date of publication. Accession numbers are listed in the key resources table. All original code will be deposited on GitHub and made publicly available on the date of publication. Any additional information required to reanalyze the data reported in this paper is available from the lead contact upon request.

### Additional resources

To facilitate the exploration of the data presented in this study, we have created the Cell Cycle database (CCdb) (Supplementary Document 2). The CCdb is a repository of proteins and phosphorylation sites exhibiting cell cycle-dependent oscillation. The data is enriched with information on protein, phosphorylation site and mRNA abundance, protein localisation, protein complexes, cancer dependency, and interaction interfaces. CCdb is available as an interactive web server at http://slim.icr.ac.uk/cell_cycle/.

### Software and server implementation

All data processing was performed in Python and R. The CCdb web server interface is written in JavaScript using the React framework. The server side is written as a Python FastAPI Web Framework. Data is stored in a PostgreSQL database. A detailed description of CCdb usage and output is provided on the help page (http://slim.icr.ac.uk/cell_cycle/blog?blog_id=ccdb_help). CCdb has been successfully tested on all major modern browsers.

## Acknowledgements

We thank Luca Cirillo for critical feedback on the manuscript, Eleanor Wendy Trotter for expert assistance on palbociclib synchronisation and the cell cycle community for their testing and suggestions for the CCdb resource. We thank Anna Santamaria and Erich Nigg for sharing the FAM83D antibody. We thank the Proteomics, Flow Cytometry, and Light Microscopy and Confocal Microscopy facilities at the ICR. This work was funded by a Cancer Research UK Senior Cancer Research Fellowship to C.R., I.T., N.E.D (C68484/A28159); and J.V and J.M (RCCSCF-Nov22/100001).

## Supplementary material

### Documents

**Document S1**: Supplementary document containing additional information on the processing of the cell cycle proteomics and phosphoproteomics datasets.

**Document S2**: Description of the Cell Cycle database (CCdb).**()Tables:**

### Tables

**Table S1**: Reviewed proteins from the UniProt Resource annotated with proteins detected in this study and other cell cycle proteomics datasets. Cyclins and proteins associated with the mitotic cell cycle process Gene Ontology (GO) term (GO:1903047) in publicly available datasets are reported.

**Table S2:** List of proteins and phosphorylation events detected in the Time Course dataset. Protein and phosphorylation abundance values are grouped by normalisation method. Statistical metrics and other functional annotations are reported. Phosphorylation sites are annotated with structural, evolutionary, functional, genomic and proteomics data.

**Table S3:** Gene Ontology (GO) term (“cell cycle” GO:0007049) enrichment analysis on different fold change ranges used to define the cutoffs for Cell Cycle-dependent (CCD) proteins in the Time Course and Mitotic Exit datasets.

**Table S4:** Gene Ontology (GO) term enrichment analysis on proteins and phosphorylation events significantly oscillating in the Time Course dataset categorised based on Biological Process, Molecular Function, Cellular Compartment and Reactome Pathway.

**Table S5:** List of proteins and phosphorylation events detected in the Mitotic Exit dataset. Protein and phosphorylation abundance values are grouped by normalisation method. Statistical metrics and other functional annotations are reported. Phosphorylation sites are annotated with structural, evolutionary, functional, genomic and proteomics data.

**Table S6:** Gene Ontology (GO) term enrichment analysis on proteins and phosphorylation events significantly enriched in the Mitotic Exit dataset categorised based on Biological Process, Molecular Function, Cellular Compartment and Reactome Pathway.

**Table S7:** Gene Ontology (GO) term enrichment analysis of proteins found in clusters shown in Figure 3F categorised based on Biological Process, Molecular Function, Cellular Compartment and Reactome Pathway.

**Table S8:** Set of Cell Cycle-dependent (CCD) proteins and phosphorylation sites. Protein and phosphorylation abundance values are grouped by normalisation method. Statistical metrics and other functional annotations are reported. Phosphorylation sites are annotated with structural, evolutionary, functional, genomic and proteomics data. Other studies defining CCD proteins and phosphorylation sites are indicated. Proteins detected in other cell cycle cell cycle datasets together with CCD proteins defined in other studies.

**Table S9:** Gene Ontology (GO) term enrichment analysis of Cell Cycle-dependent (CCD) proteins and phosphorylation events categorised based on Biological Process, Molecular Function, Cellular Compartment and Reactome Pathway.

**Table S10:** Localisation and Protein Complex enrichment analysis shown in Figure 4B. The localisation analysis table contains the localisation GO terms, the number of the proteins assigned with the respective term along with the curve fold change mean, median and standard deviation calculated from the curve fold changes of the individual proteins for a particular localisation GO term. The Protein Complexes table contains the proteins involved in the complexes along with the curve fold change mean, median and standard deviation calculated from the curve fold changes of the individual proteins involved in the complex.

**Table S11:** Proteins and transcripts identified in the Mahdessian *et al* ^15^ annotated with Cell Cycle-dependent (CCD) proteins detected in this study.

**Table S12:** Phosphorylation sites localised at protein interfaces. Statistical metrics and other functional annotations are reported.

**Table S13:** Cycle-dependent (CCD) proteins known and predicted to contain degradation motifs. Prediction confidence values and proteins containing unstable peptides ^48^ are reported.

### Supplementary Figures

**Supplementary Figure 1.**
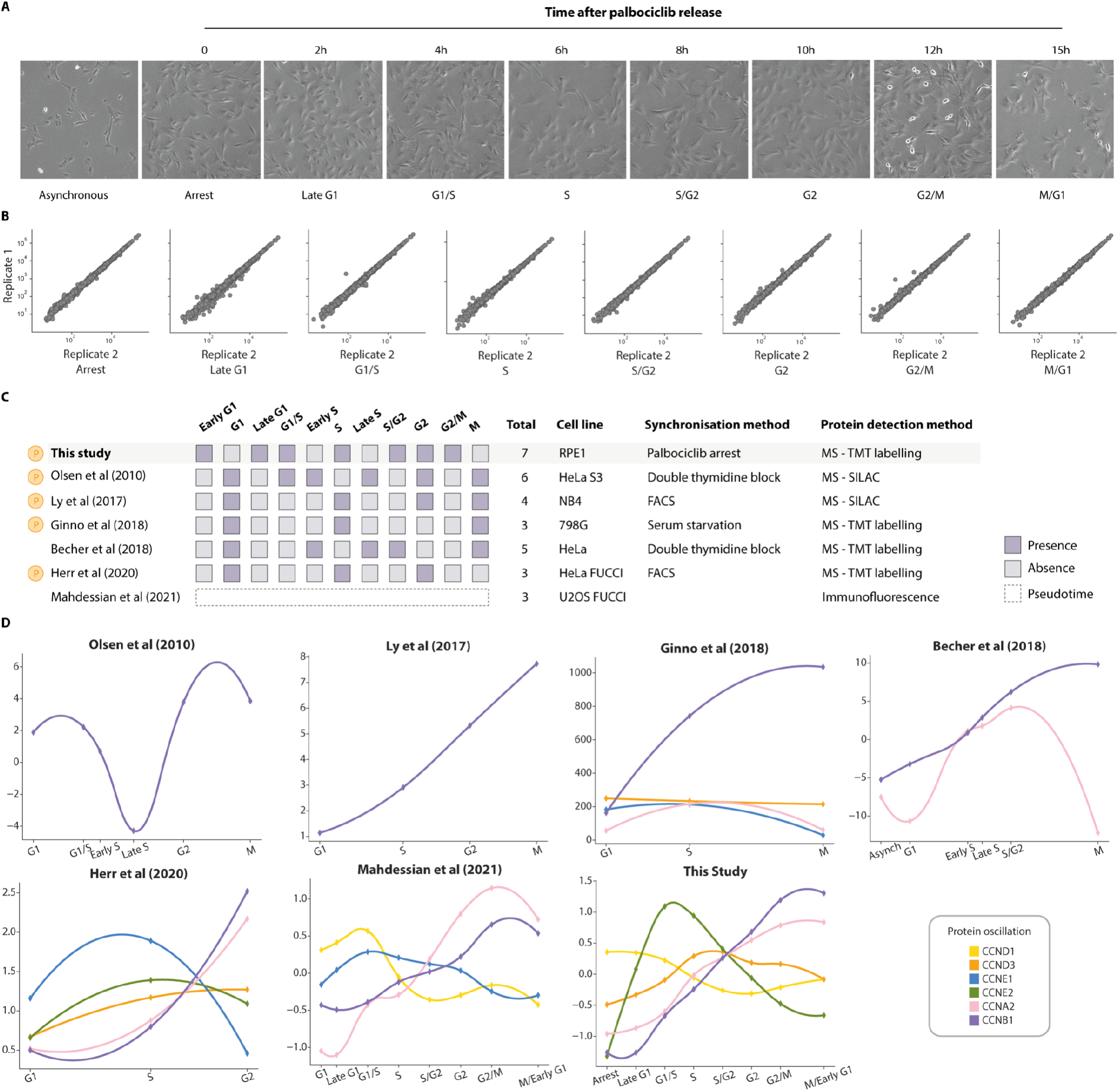
(A) Microscopy images of RPE-1 cells synchronised with palbociclib as shown in Figure 1A. (B) Multi scatter plot showing protein abundance correlation between biological duplicates. (C) Schematic representation of cell cycle phases collected in previously published cell cycle datasets. The total number of cell cycle phases collected, cell lines and methods used are shown. In the Mahdessian dataset instead of time points pseudotemporal position of each cell in interphase was measured using the FUCCI system. (D) Profile plots showing cyclin log_2_ mean normalised protein abundance oscillation during the cell cycle detected in our study compared to previously published cell cycle datasets.

**Supplementary Figure 2.**
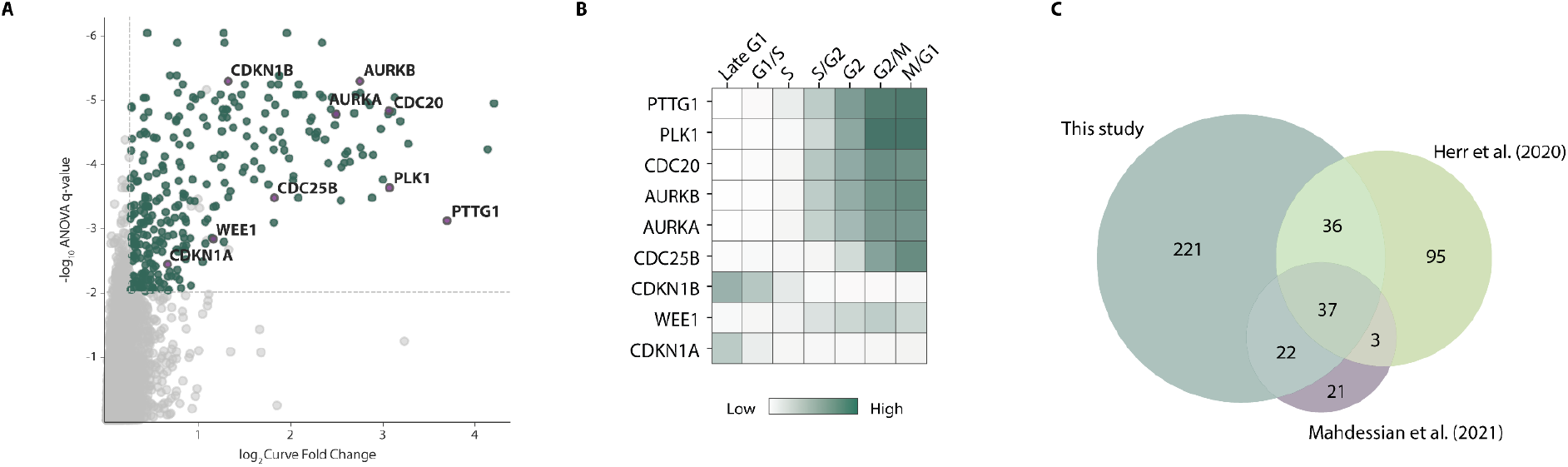
(A) Scatter plot showing well-known cell cycle-regulated proteins detected to oscillate in our proteomics analysis. The curve fold change (oscillation score) was plotted against the q-values from the ANOVA test. Significantly oscillating proteins are shown in green and non-significant in grey. Dotted lines represent cut-offs used to define oscillating proteins (ANOVA q-values ≤ 0.01, curve fold change ≥ 1.2). Well-known cell cycle-regulated proteins are highlighted. (B) Heatmap showing log_2_ mean normalised protein abundance changes of the cell cycle markers highlighted in (A). C) Venn diagram showing the overlap of oscillating phosphoproteins detected in the Time Course dataset with those detected in other cell cycle studies.

**Supplementary Figure 3.**
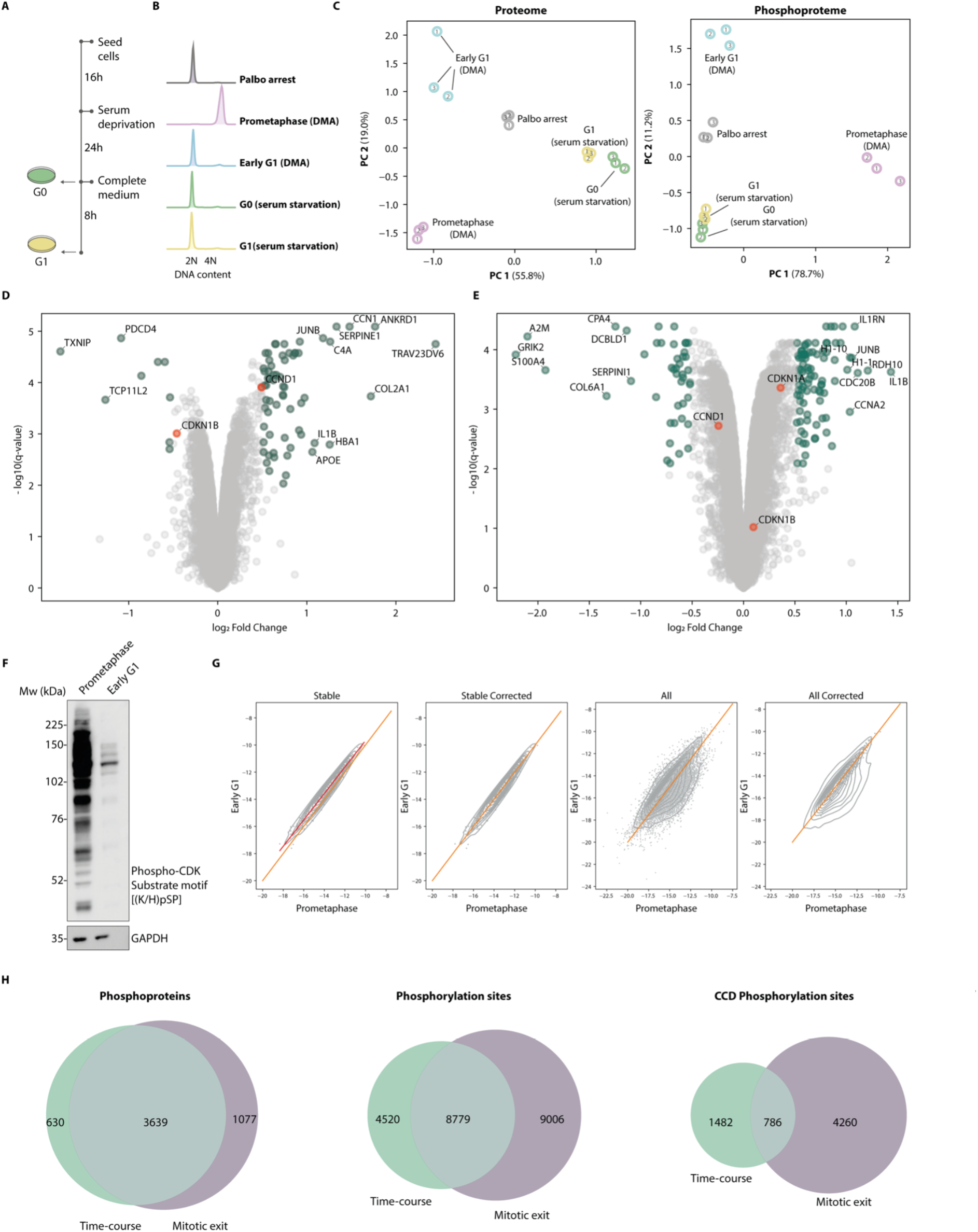
(A) Experiments workflow of cell synchronisation coupled to MS analysis. For serum starvation synchronisation, cells were incubated with serum-free medium for 24h and subsequently released in a complete medium. (B) Flow cytometry analysis assessing DNA content at each time point. (C) Principal Component Analysis of the raw abundances for the proteome and the phosphoproteome. (D) Volcano plot showing the fold change and q-values from the limma statistical test comparing serum starvation arrest (G0) and release (G1). Proteins significantly enriched are shown in green. G0 markers are highlighted in red and proteins with fold change values higher than |1| are shown. (E) Volcano plot showing the fold change and q-values from limma statistical test comparing serum starvation release (G1) and palbociclib arrest. Proteins significantly enriched are shown in green. G1 protein markers are highlighted in red and proteins with fold change values higher than |1| are shown. (F) Western blot of phospho-CDK substrate motif [(K/L)pSP] in prometaphase and early G1 samples showing the difference in global phosphorylation levels. (G) Correction of the phosphorylation sites detected in the Mitotic Exit dataset based on stable phosphorylation events detected in the Time Course dataset. Correction on replicate 1 is shown as a representative example. (H) Overlap of phosphoproteins, phosphorylation sites and CCD phosphorylation sites in the Time Course and Mitotic Exit datasets.

**Supplementary Figure 4.**
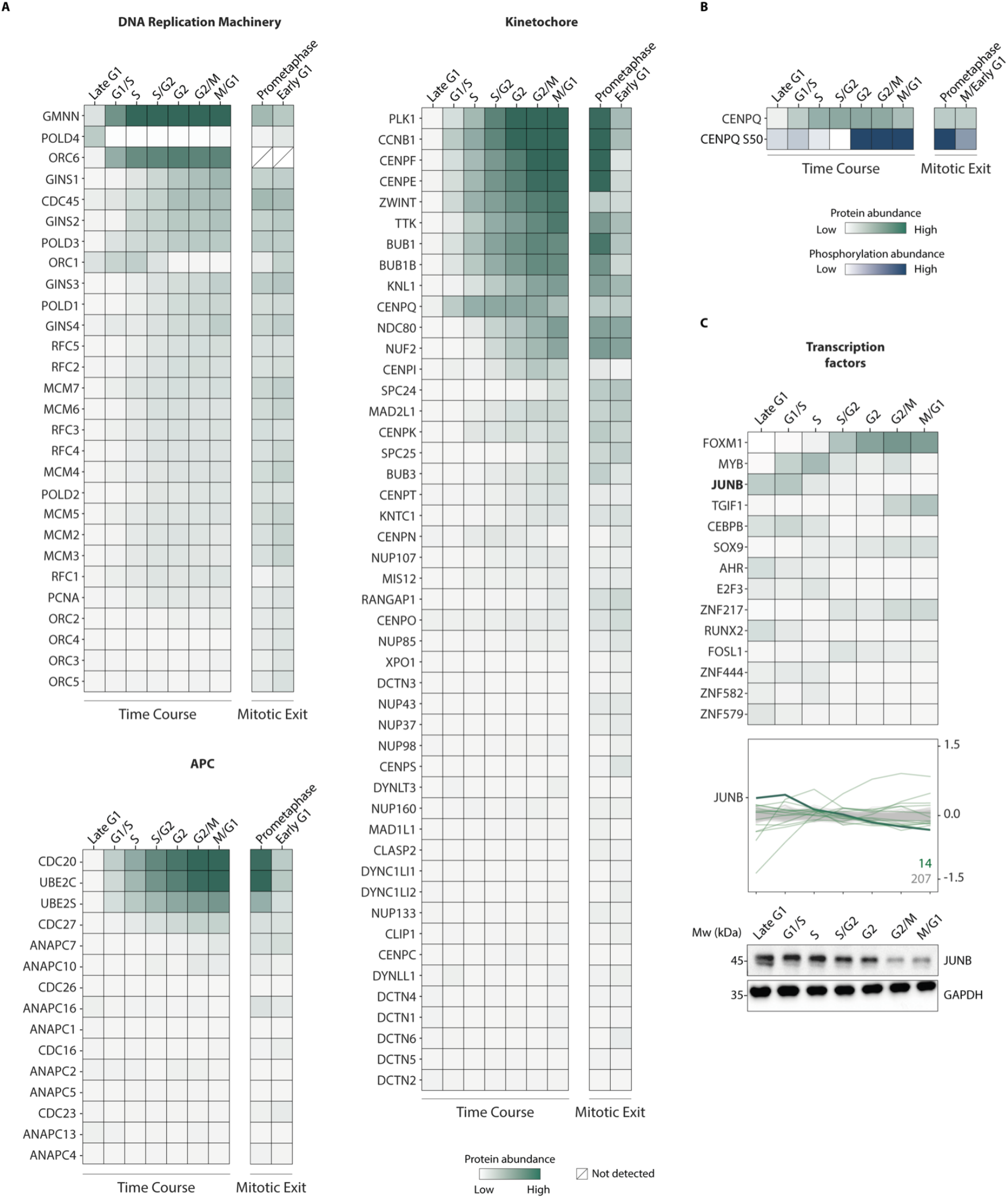
(A) Heatmaps showing log_2_ palbociclib normalised protein oscillation profiles of proteins localised at the DNA replication machinery, Anaphase Promoting Complex/Cyclosome and kinetochore. (B) Heatmaps showing the CENPQ log_2_ palbociclib normalised protein (green) and pS50 (blue) abundance changes during the cell cycle detected by MS during cell cycle progression in both Time Course and Mitotic Exit datasets. (C) Heatmap of the log_2_ mean normalised abundance for transcription factors detected in the Time Course dataset and profile plots showing the most significantly oscillating proteins. Numbers of oscillating (green) and stable (grey) proteins are shown and proteins validated by western blot are in bold.

**Supplementary Figure 5.**
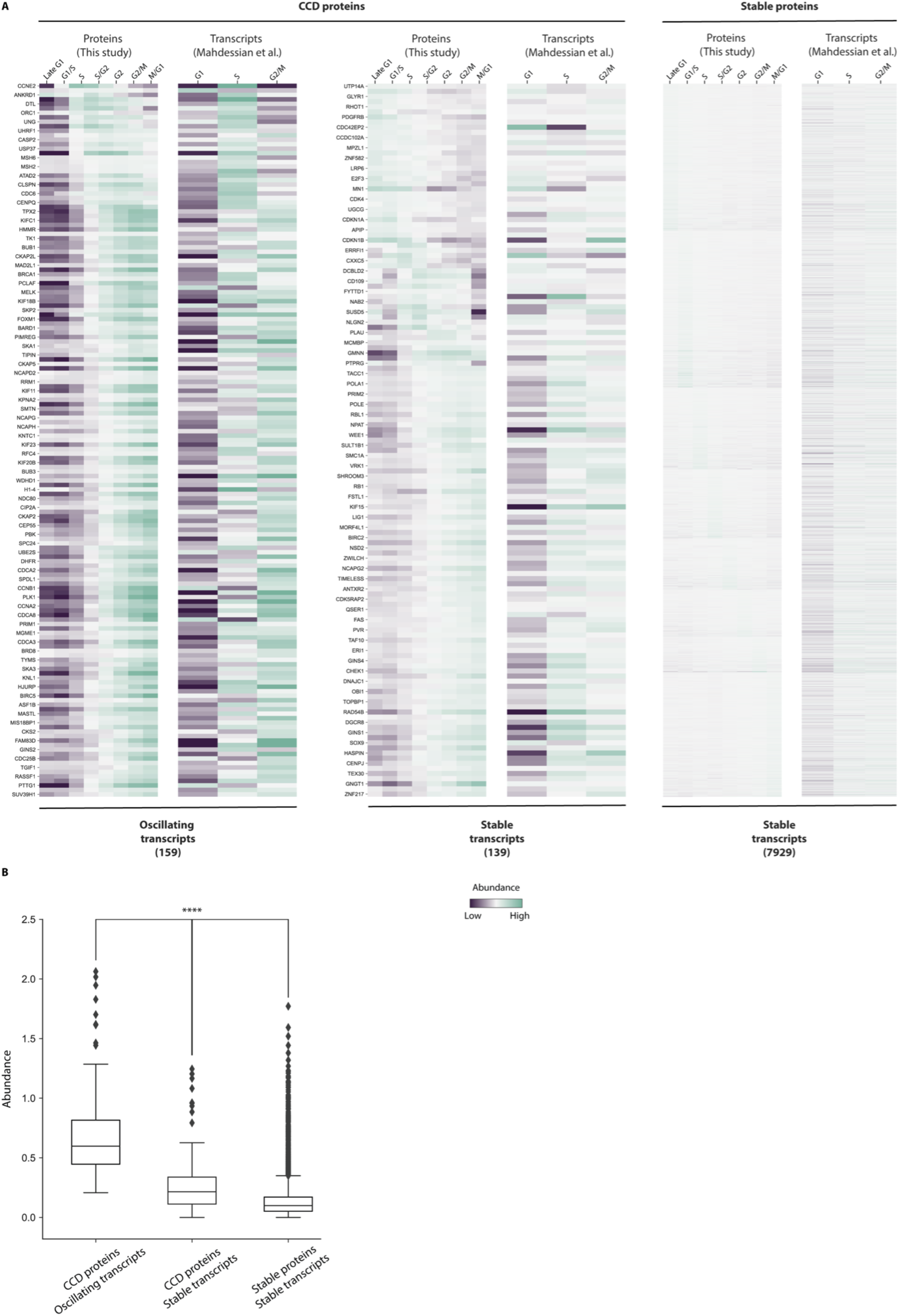
(A) Heatmap of the log_2_ mean normalised protein and transcript abundance from this study (Time Course dataset) and Mahdessian *et al* ^15^ respectively. Protein and transcript abundance are categorised into CCD and stable groups using this dataset for proteins and Mahdessian *et al* ^15^ for transcripts. (B) Boxplots of the maximum abundance for each transcript in the categories shown in (A). All categories are significantly different from each other (**** Mann-Whitney p-value < 10-17). The transcripts belonging to the first category (in which both transcript and protein are CCD) are the most abundant.

**Supplementary Figure 6.**
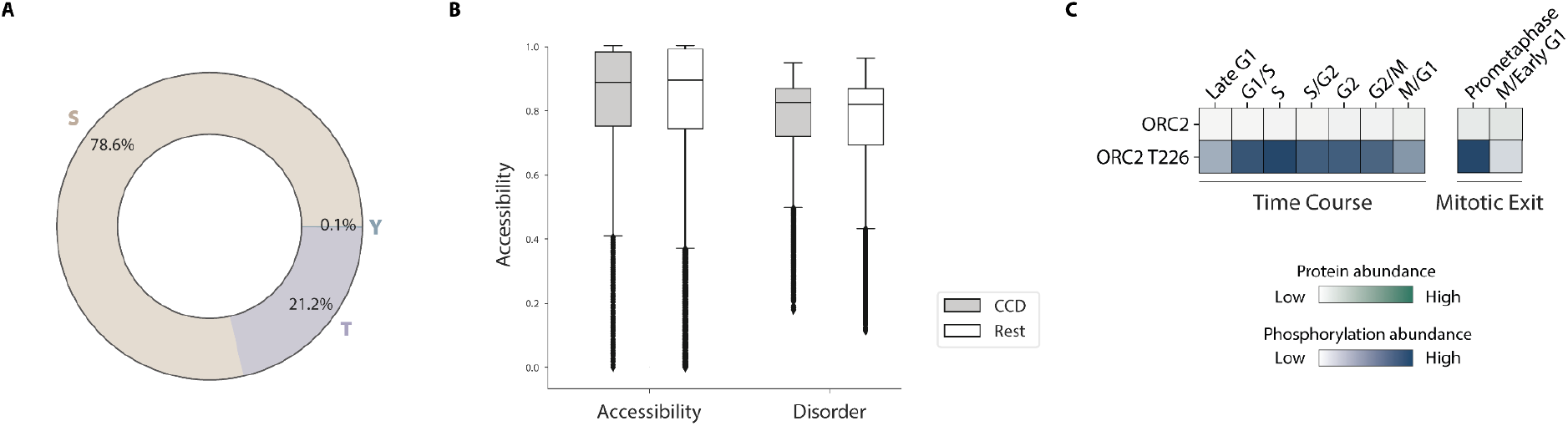
(A) Percentages of phosphorylated serine, threonine and tyrosine residues detected in the CCD phosphoproteome dataset. (B) AlphaFold2 Accessibility and Disorder of the CCD phosphorylation sites in comparison to the rest. (C) Heatmap showing the ORC2 log_2_ mean normalised protein (green) and pThr226 (blue) changes detected by MS during cell cycle progression in both Time Course and Mitotic Exit datasets.

**Supplementary Figure 7.**
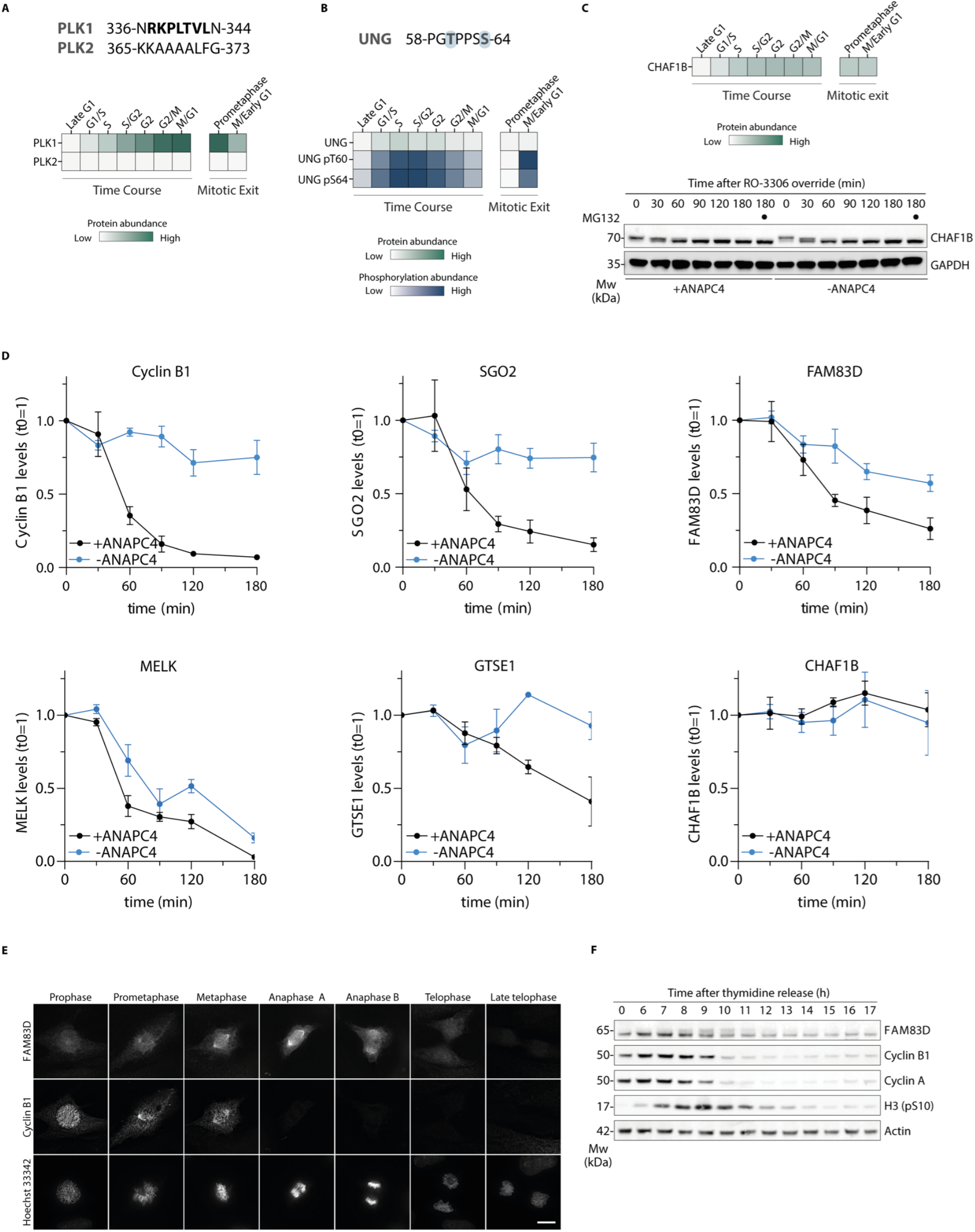
(A) Sequence alignment of PLK1 and PLK2 showing lack of D box motif in PLK2. PLK1 and PLK2 log_2_ palbociclib normalised protein changes during cell cycle progression detected by MS in both Time Course and Mitotic Exit datasets are shown in the heatmap (bottom). (B) The amino acid sequence of phosphopeptide degron in UNG. UNG protein (green) and its phosphorylation (blue) log_2_ palbociclib normalised changes during cell cycle progression detected by MS in both Time Course and Mitotic Exit datasets are shown in the heatmap (bottom). (C) Heatmap showing CHAF1B log_2_ palbociclib normalised protein changes during cell cycle progression detected by MS in both Time Course and Mitotic Exit datasets (top). Western blot analysis of CHAF1B degradation in Hela cells during the override of a mitotic checkpoint arrest by the CDK1 inhibitor RO-3306 in the presence and absence of ANAPC4 (bottom). (D) Quantification of western blots from Figure 6D (n=3). (E) Immunostaining of RPE-1 cells with anti-cyclin B1 and anti-FAM38D antibodies. Scale bar = 10 µm (n=2). (F) Western blot analysis FAM38D protein levels in Hela Kyoto cells released from a thymidine-block in G1/S phase (n=2).

## Notes

https://slim.icr.ac.uk/cell_cycle/

### Competing Interest Statement

The authors have declared no competing interest.

