## Supplementary Table for "High resolution profiling of cell cycle-dependent protein and phosphorylation abundance changes in non-transformed cells": Supplementary Document 1.docx

### ***Normalisation***

The initial protein and phosphorylation site abundances from the MS data processing, here defined as *raw protein abundances*, for all sets, were normalised by dividing the median intensity for each sample prior to further analysis to account for changes in the total cell content during the cell cycle to produce the *median normalised protein abundance*. Subsequently, protein and phosphorylation abundances were normalised for each dataset to allow specific analyses as follow:

Time Course Dataset:

The Time Course dataset contains protein and phospho-peptide abundances for 16 samples (two replicates across eight time-points). Two metrics were used in the study to analyse the Time Course data: the *log2 mean* and *0-max* metrics. The *Log2 mean* metric calculates the relative abundance of a time point against the mean abundance across all time points. For each protein or phosphosite, the median normalised abundances of the 16 samples were log2 transformed and the abundances were subtracted from the mean log2 abundances of the 16 samples.

${Na}_{rt}$ = log2($a_{rt}$) - $a_{mean}$

***Equation 1:*** *The* *log2-mean abundance normalisation.* ${Na}_{rt}$ *is the log2-mean abundance normalisation for a given time point (t) and replicate (r),* $a_{rt}$ *is the median normalised protein abundance for a given time point (t) and replicate (r), and* $a_{mean}$ *is the mean of the log2 normalised abundances for a protein or phosphorylation sites for both replicates.*

The *0-max* metric calculates the abundance of a time point relative to the maximum abundance across all time points. For each protein and phosphorylation site, the 16 median normalised abundances were divided by the maximum median normalised abundances. A related *min-max* metric was also calculated where the minimum median normalised abundance is subtracted before performing the *0-max* normalisation resulting in an abundance profile where the minimum value is 0 and maximum value is 1.

${Na}_{tr}$ = $\frac{a_{tr}}{a_{max}}$

***Equation 2:*** *The* *0 - max abundance normalisation.* ${Na}_{tr}$ *is the*  *0 - max abundance normalisation for a given time point (t) and replicate (r),* $a_{rt}$ *is the median normalised protein abundance for a given time point (t) and replicate (r) and* $a_{max}$ *is the maximum median normalised abundance for the protein or phosphorylation sites.*

Mitotic Exit Dataset:

The Mitotic Exit dataset contains protein and phospho-peptide abundances for 15 samples (three replicates across five time points). The *log^2^ sum* metric was used in the study to analyse the Mitotic Exit data. For each protein and phosphorylation site, the *log^2^ sum* is calculated as the log^2^ of the raw abundances divided by the sum of all the raw abundances for a specific time point and replicate.

${Na}_{rt}$ = log2($\frac{a_{rt}}{a_{col Sum}}$)

***Equation 3:*** The *log^2^ sum abundance normalisation. Where* ${Na}_{t}$ *is the log^2^ sum protein abundance normalisation for a given time point (t) and replicate (r),* $a_{rt}$ *is the raw abundance for a given time point (t) and replicate (r), and* $a_{col Sum}$ *is the abundance of all proteins or phosphorylation sites in a given time point (t) and replicate (r).*

Merging Time Course and Mitotic Exit Datasets:

The *log2 Palbociclib* metric is used to merge the Time Course and Mitotic Exit datasets. Both sets include Palbociclib time points allowing the data to be integrated. For each protein and phosphorylation site, the median normalised abundances for each replicate and each time-point were divided by the mean Palbociclib abundance for the time-point and then log2 transformed to scale all data points relative to the Palbociclib data.

${Na}_{rt}$ = log2($\frac{a_{rt}}{a_{r,palbo}}$)

***Equation 4:*** *The log^2^ Palbociclib* *abundance normalisation. Where* ${Na}_{rt}$ *is the log^2^ Palbociclib protein abundance normalisation for a given time point (t) from a given replicate (r),* $a_{rt}$ *is the median normalised protein abundance for a given time point (t) and replicate (r), and* $a_{r,palbo}$ *is the median normalised protein abundance for the time point Palbociclib for a given replicate (r).*

***Correction of the Mitotic Exit phosphorylation sites abundances***

The total intensity of phosphopeptides in the DMA Arrest was double compared to the other conditions. To correct for this the phosphorylation sites were split into two categories: stable, based on the “Stable Sites” definition of the time course dataset and the remaining sites. The "stable sites" category was subsequently refined by removing the outliers in the top by 10% of lower and upper bounds of the stable sites distribution, resulting in a more compact and precisely defined set of stable sites. A linear model was fitted on the DMA Arrest and DMA Release time points of the stable sites set using the *linear_model.LinearRegression* function from the *sklearn* library and the slope and intercept were calculated. The corrected DMA Arrest time point per replicate for each phosphorylation was then calculated.

***Accessibility calculation***

Accessibility is calculated from DSSP residue accessible surface area (ASA) of AlphaFold2 models [PMID: 36344848]. Accessibility is the surface accessibility score derived from AlphaFold2 models for the single phosphorylation site normalised by the maximum possible accessibility for that residue (*Equation 7*).

${SA}_{p} =\frac{A_{p}}{M_{p}}$

**Equation 7: Surface Accessibility score for specific position p in protein.**

${SA}_{p}$ is the square Ångstrom surface area of a residue at position *p* in the structure that can be accessed by a water molecule. M is the theoretical maximum Ångstrom surface area of the amino acid at position *p.* The values for M are A:124, C:94, D:154, E:187, F:221, G:89, H:201, I:193, K:214, L:199, M:216, N:161, P:130, Q:192, R:244, S:113, T:151, V:169, W:264 and Y:237. M values were calculated for 5-residue peptides with a central query amino acid flanked by two glycines (e.g. GGXGG).
