## Supplementary Table for "High resolution profiling of cell cycle-dependent protein and phosphorylation abundance changes in non-transformed cells": Supplementary Document 2.docx

#### **Description of the Cell Cycle Database (CCdb)**

#### ***Data sources***

**In-house Datasets**

The main datasets stored in the CCdb are the data produced during the current study. These data consist of three distinct analyses. The datasets were categorised based on the synchronisation method used to produce them: **(i)** a cell cycle Time Course dataset, obtained by palbociclib (CDK4/6 Inhibition) induced arrest, covering seven cell cycle stages from late G1 to M/G1 (Figure 1), (ii) a Mitotic Exit dataset, obtained by Dimethylenastron (DMA) (Eg5 inhibition) induced arrest, covering prometaphase and Early G1 (Figure 2) and (iii) a Serum Starvation dataset, obtained by serum starvation-induced arrest, covering G0 and G1 (Figure 3). These datasets are described in more detail in the methods section of the paper. However, an overview of the dataset categories is the following:

**(i)	Time Course dataset:**

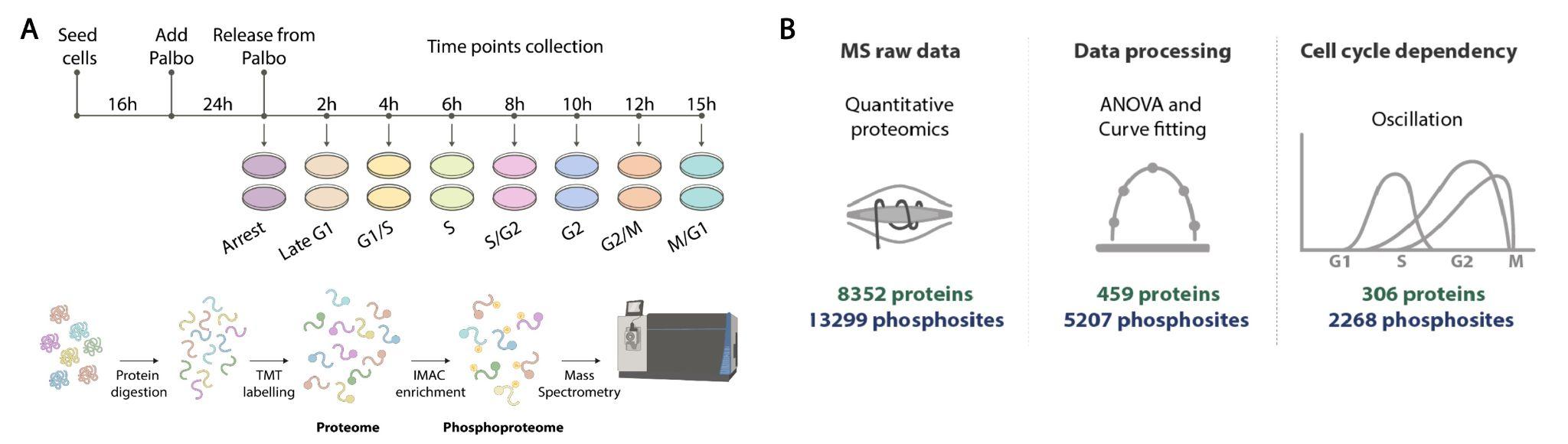

**Figure 1.** *Overview of the Time Course experimental design and results summary. (A) General experiments workflow of cell synchronisation coupled to Mass Spectrometry (MS) analysis. Cells were arrested in late G1 with palbociclib (Palbo) and time points corresponding to different cell cycle phases were collected upon release from the inhibitor. Sample aliquots were fixed, and DNA was stained with propidium iodide for flow cytometry. For MS analysis, cells were lysed, and protein digested with trypsin, followed by TMT labelling and phosphopeptides enrichment. (B) Simplified data analysis workflow used to define oscillating proteins and phosphosites.*

**(ii)	Mitotic Exit dataset:**

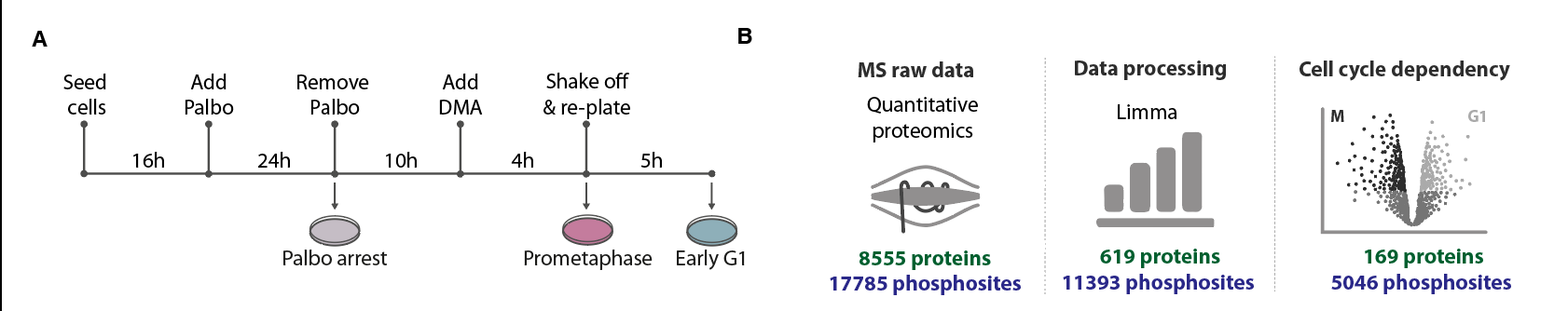

**Figure 2.** *Overview of the Mitotic Exit experimental design and results summary. (A) Experimental workflow of cell synchronisation coupled to MS analysis. Cells synchronised in G2 with palbociclib-induced arrest were incubated with DMA for 4h. Prometaphase-arrested cells were harvested by mitotic shake-off and released in a fresh medium. Upon 5h release, early G1 cells were collected. (B) Simplified data analysis workflow used to define proteins and phosphorylation events significantly changing in DMA arrest (prometaphase) and DMA release (early G1).*

**(iii)	Serum Starvation dataset:**

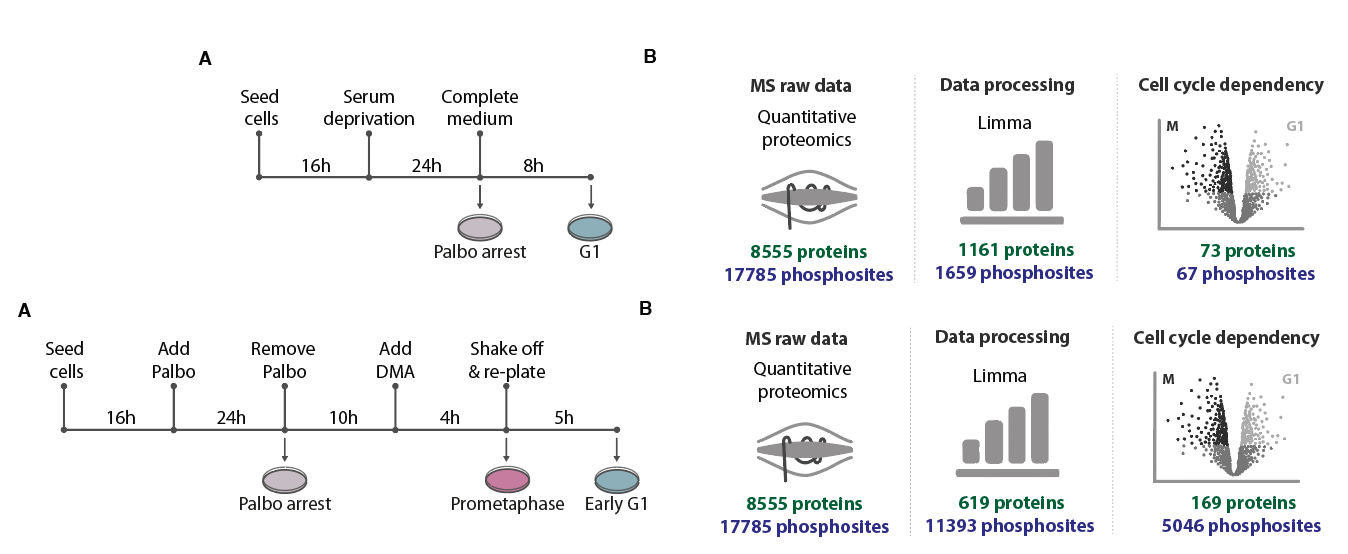

**Figure 3.** *Overview of the Serum Starvation experimental design and results summary.* *(A) Experimental workflow of cell synchronisation coupled to MS analysis. To collect the G0 phase, cells were serum-deprived for 24h and subsequently cultured in a complete medium. The G1 phase was collected after 8h release from serum deprivation. (B) Simplified data analysis workflow used to define proteins and phosphorylation events significantly changing in serum starvation arrest (G0) and serum starvation release (G1).*

#### ***CCD Datasets***

The resource identifies proteins and phosphorylation sites exhibiting cell cycle-dependent oscillation by combining these MS abundances with statistical metrics. The serum starvation-induced arrest revealed limited protein or phosphorylation abundance changes, therefore, we defined the core cell cycle-dependent (CCD) set for proteins and phosphorylation sites based on the Time Course and Mitotic Exit datasets (Figure 4).

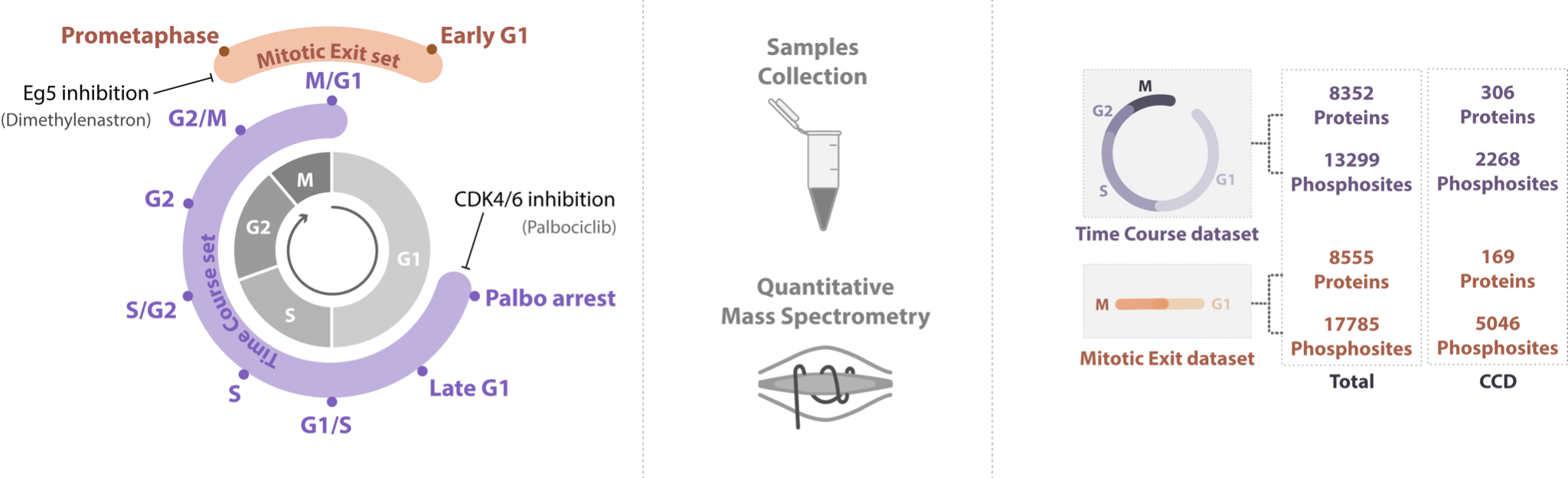
**Figure 4.** *Overview of the time points collected for the CCdb datasets, a simplified overview of the experimental design, and final results.*

#### ***External Datasets***

The CCdb resource has been extensively enriched by the addition of complementary datasets from numerous resources. The following datasets have been parsed and integrated with the previously described in-house datasets.

##### ***Proteomics Datasets***

External proteomics datasets were retrieved from the supplementary material of the source publications, processed and mapped to UniProt identifiers.

| **Name** | **Quantification Method** | **Cell Line** | **Synchronisation Method** | **Cell Cycle Phases** | **CCD** | **PMID** |
| --- | --- | --- | --- | --- | --- | --- |
| Mahdessian et al. (2021) | Immunofluorescence | U2OS | Fucci system expression, single-cell imaging | Pseudotime | True | 33627808 |
| Herr et al. (2020) | TMT labelling | Hela | Fucci system expression, FACS sorting | G1, S, G2/M | True | 32051232 |
| Ginno et al., (2018) | TMT labelling | T98G | Serum starvation 72h, Nocodazole | G1, S, M | False | 30279501 |
| Becher et al, (2018) | TMT labelling to (protein thermal stability and solubility) | Hela Kyoto | Double thymidine block, Nocodazole | G1/S, Early S, Late S, S/G2, M, Early G1 | False | 29706546 |
| Ly et al., (2017) | SILAC | NB4 | FACS sorting | G1, S, G2, M | False | 29052541 |
| Olsen et al. (2010) | SILAC (in-gel digestion) | HeLa S3 | Double thymidine block, Nocodazole | G1/S, Early S, Late S, S/G2, M, G1 | False | 20068231 |

**Table 1.** *Publicly available proteomics datasets from previously published high throughput proteomics experiments of cell cycle protein abundance. The protein quantification method, cell lines, and synchronisation methods used together with time points collected are reported. Datasets defining CCD proteins are indicated as “True”.*

##### ***Phosphoproteomics datasets***

External phosphoproteomics datasets were retrieved from the supplementary material of the source publications, processed and mapped to UniProt identifiers and positions.

| **Name** | **Methods** | **Cell Line** | **Synchronisation Method** | **Cell Cycle Phases** | **CCD** | **PMID** |
| --- | --- | --- | --- | --- | --- | --- |
| Herr et al. (2020) | TMT labelling and TiO2 phospho-enrichment | Hela | Fucci system expression, FACS sorting | G1, S, G2/M | False | 32051232 |
| Ginno et al., (2018) | TMT labelling | T98G | Serum starvation 72h, Nocodazole | G1, S, M | False | 30279501 |
| Ly et al., (2017) | TMT labelling and TiO2-IMAC phospho-enrichment | NB4 | FACS sorting | G1, S, G2, M | False | 29052541 |
| Olsen et al. (2010) | SILAC (in-gel digestion) and TiO2-IMAC phospho-enrichment | Hela S3 | Double thymidine block, Nocodazole | G1/S, Early S, Late S, S/G2, M, G1 | True | 20068231 |
| McCloy et al. (2010) | SILAC and TiO2 phospho-enrichment | Hela | Double thymidine block, Nocodazole | - | True | 26055452 |

**Table 2.** *External, publicly available datasets from previously published high throughput phosphoproteomics experiments of cell cycle phosphorylation site abundance. Phosphorylation sites abundance quantification method, cell lines, and synchronisation methods used together with time points collected are reported. Datasets defining CCD phosphorylation sites are indicated as “True”.*

##### ***Transcriptomics datasets***

External transcriptomics datasets were retrieved from the supplementary material of the source publications, processed and mapped to UniProt identifiers.

| **Name** | **Methods** | **Cell Line** | **Synchronisation**  **Method** | **Cell Cycle Phases** | **CCD** | **PMID** |
| --- | --- | --- | --- | --- | --- | --- |
| Mahdessian et al. (2021) | Smart-Seq2 | U2OS | Fucci system expression, single-cell imaging | Pseudotime | True | 33627808 |
| Herr et al. (2020) | TruSeq RNA | Hela | Fucci system expression, FACS sorting | G1, S, G2/M | False | 32051232 |
| Ficher et al. (2020) | Meta-analysis | HCT116 | Fucci system expression, FACS sorting | G1/S, G2/M | True | 27280975 |
| Giotti et al., (2017) | Meta-analysis | - | - | - | True | 28056781 |
| Grant et al., (2013) | Microarray hybridization,  ChIP-seq | U2OS | Double thymidine block, Nocodazole | G1/S, S, G2, G2/M, M/G1 | True | 24109597 |
| Peña-Diaz et al., (2013). | Microarray hybridization | HaCaT | Double thymidine block | G1/S, S, G2, G2/M, M/G1 | True | 23325852 |
| Bar-Joseph et al., (2008) | Microarray hybridization + Time-Lapse Cinematography | NHDF | Double thymidine block, serum starvation, FACS | G1, S, G2/M | True | 18195366 |
| Whitfield et al., (2002) | Microarray Hybridization | Hela S3 | Double thymidine block, Nocodazole | G1/S, S, G2, G2/M | True | 12058064 |

**Table 3.** *External, publicly available datasets from previously published high throughput transcriptomics experiments of cell cycle RNA abundance. mRNA quantification method, cell lines, and synchronisation methods used together with time points collected are reported. Datasets defining CCD mRNA are indicated as “True”.*

#### ***General protein feature data***

Additionally to the cell cycle-focused datasets above, data from several protein-centric databases have been parsed and integrated into CCdb to allow protein and phosphorylation information to be enriched with relevant complementary data (Table 4). This includes information on: interactions from IntAct, CORUM and Complex Portal to investigate the interaction network of the proteins; Protein essentiality and cancer-links from DepMap and DRIVE to define the therapeutic relevance of the proteins; Protein architecture from Pfam, PDB and AlphaFold to understand phosphorylation site context; Phosphorylation site from Phospho.ELM, PhosphoSitePlus and Ochoa et al [PMID: 31819260] to define the confidence and novelty of the observed phosphorylation site.

| **Name** | **Description** | **PMID** |
| --- | --- | --- |
| UniProt | Protein accessions, names, sequences, families, UniRef clusters and feature annotations. | 25348405 |
| ELM | Manually curated linear motifs. | 34718738 |
| Pfam | Functional regions and binding domains. | 24288371 |
| Phospho.ELM | Experimentally verified phosphorylation sites. | 21062810 |
| PhosphoSitePlus | Phosphorylation, ubiquitination, acetylation and methylation sites. | 22135298 |
| Ochoa et al. - PTMs | Phosphoproteomic dataset. | 31819260 |
| PDB | Experimentally resolved protein tertiary structures. | 10592235 |
| AlphaFold DB | 3D model predictions of protein structures. | 34265844 |
| DepMap | Gene dependencies in hundreds of cancer cell lines. | 28753430 |
| Cancer Gene Census (CGC) | Catalogue of genes which contain mutations that have been causally implicated in cancer. | 30293088 |
| DRIVE cancer | Driver genes derived from a PanCancer and PanSoftware analysis. | 29625053 |
| IntAct | Experimentally validated protein-protein interactions. | 24234451 |
| CORUM | A comprehensive resource of mammalian protein complexes. | 17965090 |
| Complex Portal | Manually curated, encyclopaedic resource of macromolecular complexes from several key model organisms. | 30357405 |

**Table 4.** *Databases used and integrated into CCdb, their description and reference*.

#### ***Protein stability information***

A range of data related to protein stability and targeting the ubiquitin-proteasome system (UPS) was mapped to the CCdb data to link cell cycle-dependent abundance dynamics to degradation. A set of degrons, a class of degradation motifs that recruits components of the UPS to drive protein polyubiquitination and subsequent degradation, was collected to allow the degradation mechanisms for proteins to be annotated. Degrons were curated from three sources: the validated APC/C degrons derived from Davey *et al.* [PMID: 27716480], the ELM database [PMID: 34718738] and manual curation. These data were augmented by a set of destabilising peptides from a global peptide stability (GPS) screen of ∼470,000 28-mer peptides that identified 15,800 degron-containing peptides [PMID: 37738965]. A dataset of protein half-life defined by cycloheximide chase coupled to mass spectrometry [PMID: 34626566] was processed and each protein in the Time Course and Mitotic Exit datasets was annotated with the relative abundance mean and standard deviation for 8h for each protein. These data allow cycle-dependent abundance dynamics to be cross-validated by observed low protein stability.

| **Name** | **Description** | **PMID** |
| --- | --- | --- |
| ELM | Manually curated linear motifs. | 34718738 |
| GPS | Proteome-wide global protein stability (GPS) assay | 37738965 |
| Protein half-life | Cycloheximide chase coupled to mass spectrometry screen of protein stability | 34626566 |

**Table 5.** *External sources of protein stability-related information integrated into CCdb, their description and reference*.

#### ***Visualisation tools***

Finally, phosphorylation sites were linked to the ProViz protein visualisation software to view each site in the context of functional features and evolutionary data and allow potentially interesting observations for further investigation.

| **Name** | **Description** | **PMID** |
| --- | --- | --- |
| ProViz | Web-based visualisation tool to investigate the functional and evolutionary features of protein sequences | 27085803 |

**Table 6.** *Tool used to visualise phosphorylation site context.*

#### ***Phosphorylation sites annotation***

Phosphorylation sites were annotated with the in-house peptide annotation software PepTools [[PMID: 35044719]](https://sciwheel.com/work/citation?ids=12301587&pre=&suf=&sa=0) developed to analyse the peptide data from experimental high-throughput motif discovery methods. Peptools enriched the phosphoproteomic data with structural, evolutionary, functional, genomic and proteomics data. The main source of PTMs in PepTools is Phospho.ELM [PMID: 21062810], PhosphoSitePlus [PMID: 22135298], UniProt [PMID: 25348405] and Ochoa et al. [PMID: 31819260]. Accessibility was calculated as the surface accessibility score derived from AlphaFold2 models for the amino acids observed to be phosphorylated [PMID: 34265844]. Furthermore, a disorder score, calculated as a windowed accessibility score for a given site with a flanking region of 15 amino acids, was reported to define the regional accessibility with higher local accessibility strongly correlating with intrinsically disordered regions. Phosphorylation sites overlapping interaction interfaces were annotated based on (i) sites overlapping experimentally validated short linear motifs derived from ELM [PMD:34718738] and a set of in-house curated motifs from the MoMap database (<http://slim.icr.ac.uk/momap/>, in preparation]), (ii) sites overlapping an annotated domain or family descriptor were derived from Pfam [PMID: 24288371] and UniProt [PMID: 25348405], (iii) sites overlapping a structurally interface solved by EM, NMR or X-ray crystallography from the PDB database [PMID: 10592235]. Interface residues were determined as residues with heavy atoms within less than 6 Å distance from another protein chain in structures of protein complexes with 2 or more subunits.

#### ***Implementation***

Data processing for the CCdb was performed in Python and R and they were loaded into a PostgreSQL database. The CCdb web server interface is written in JavaScript using the React framework. The server side is written as a Python FastAPI Web Framework. A detailed description of CCdb usage and output is provided on the help page (<http://slim.icr.ac.uk/cell_cycle/blog?blog_id=ccdb_help>). CCdb has been successfully tested on all major modern browsers.

### ***General Layout***

The Cell Cycle Database can be accessed online at <http://slim.icr.ac.uk/cell_cycle/>. The home page of the CCdb web interface is shown in Figure 5. The page allows the user to browse individual proteins and phosphorylation events by selecting *Search* in the toolbar to find a specific protein (label 1). A user can also search by protein name, gene name or UniProt accession in the “Find your protein Section” of the home page. Alternatively, the complete cell-cycle dependent Time Course and Mitotic Exit datasets can be accessed through the *Proteins* tab in the toolbar (label 2). The figures from the paper can be accessed as interactive figures in the *Interactive Figures* tab in the toolbar (label 3) to explore more complex aspects of the dataset (described below).

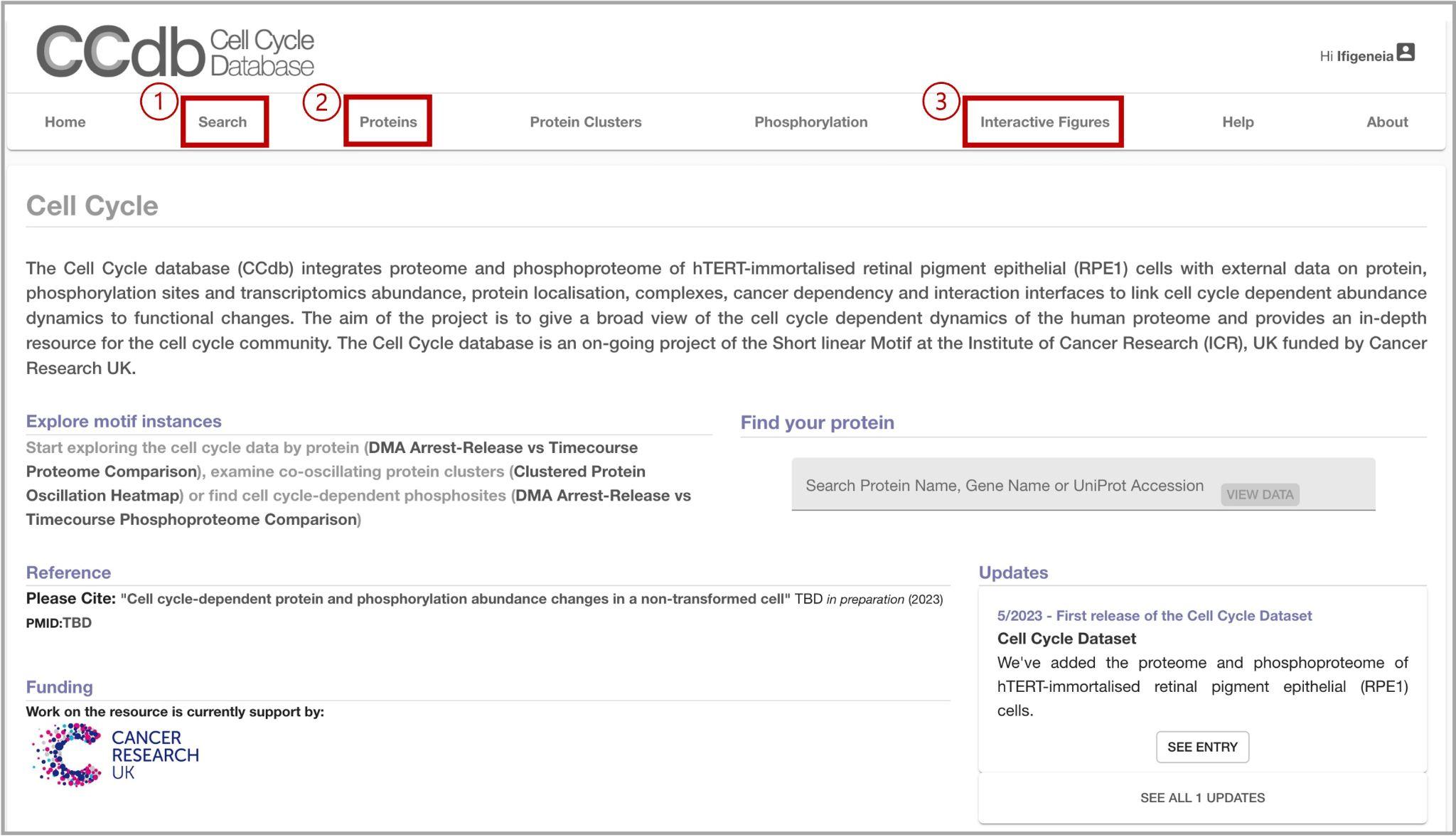

***Figure 5.*** *Overview of CCdb (Cell Cycle Database) Home page. (1) Access to the search page for the protein of interest. (2) Access to the full list of CCD proteins. (3) Access the full list of interactive Figures and their description.*

#### ***Proteins View***

A full list of the cell cycle-dependent proteins, their statistical metrics and their normalised abundances can be explored in the Proteins view (Figure 6) and accessed through the *Proteins* tab in the toolbar. Clicking on any of the protein names in the table will take the user to the individual Proteins View. The table is interactive and details on protein abundance values are shown by hovering over the heatmaps. Each column can be sorted by clicking on the table headers.

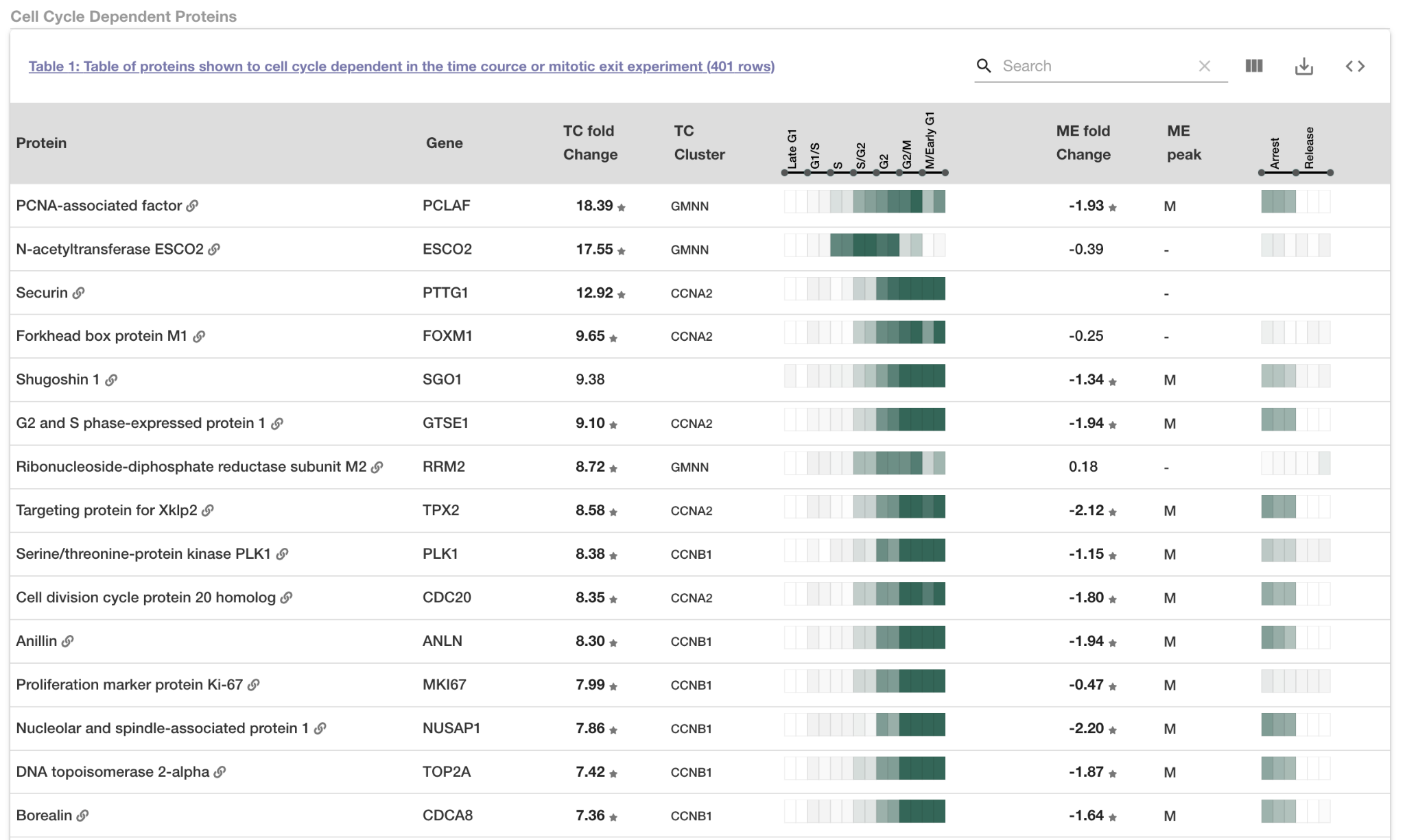

***Figure 6.*** *Screenshot of the CCD proteins table showing the top oscillating proteins in the Time Course dataset.*

#### ***Protein Instance View***

Selecting a protein in the Proteins View table or Search bar in the Search tab or homepage will take the user to the Protein Instance View (Figure 7).

##### ***Protein Abundance Overview Section***

The *protein abundance overview* section of the Protein Instance View is the overview of the protein’s cell cycle dependency (CCD) status. The section displays the CCD status for four datasets: the total combined dataset, the single Time Course, the Mitotic Exit and the Serum Starvation datasets. If the protein is cell cycle-dependent in a given dataset it is marked with a ✓ otherwise it is marked with an ✗. CCD proteins from the combined dataset have been clustered based on the similarity of their abundance profiles to known cell cycle markers (see *Interactive Figures section*). Information about the oscillation cluster is shown for CCD these proteins. For each of the single datasets, statistical metrics related to the cell cycle dependency and the peak abundance time point are displayed. If the protein is not observed in a particular screen no data is shown. The size and confidence of the protein oscillation is presented with the Curve Fold Change and ANOVA significance for the Time Course dataset, and *limma* Fold Change and adjusted p-values for the Mitotic Exit and Serum Starvation datasets. Finally, the peak phase indicates the cell cycle phase with the highest protein abundance. For a description of the different datasets see the *Data Sources* section of the Methods (Figures 1, 2 and 3).

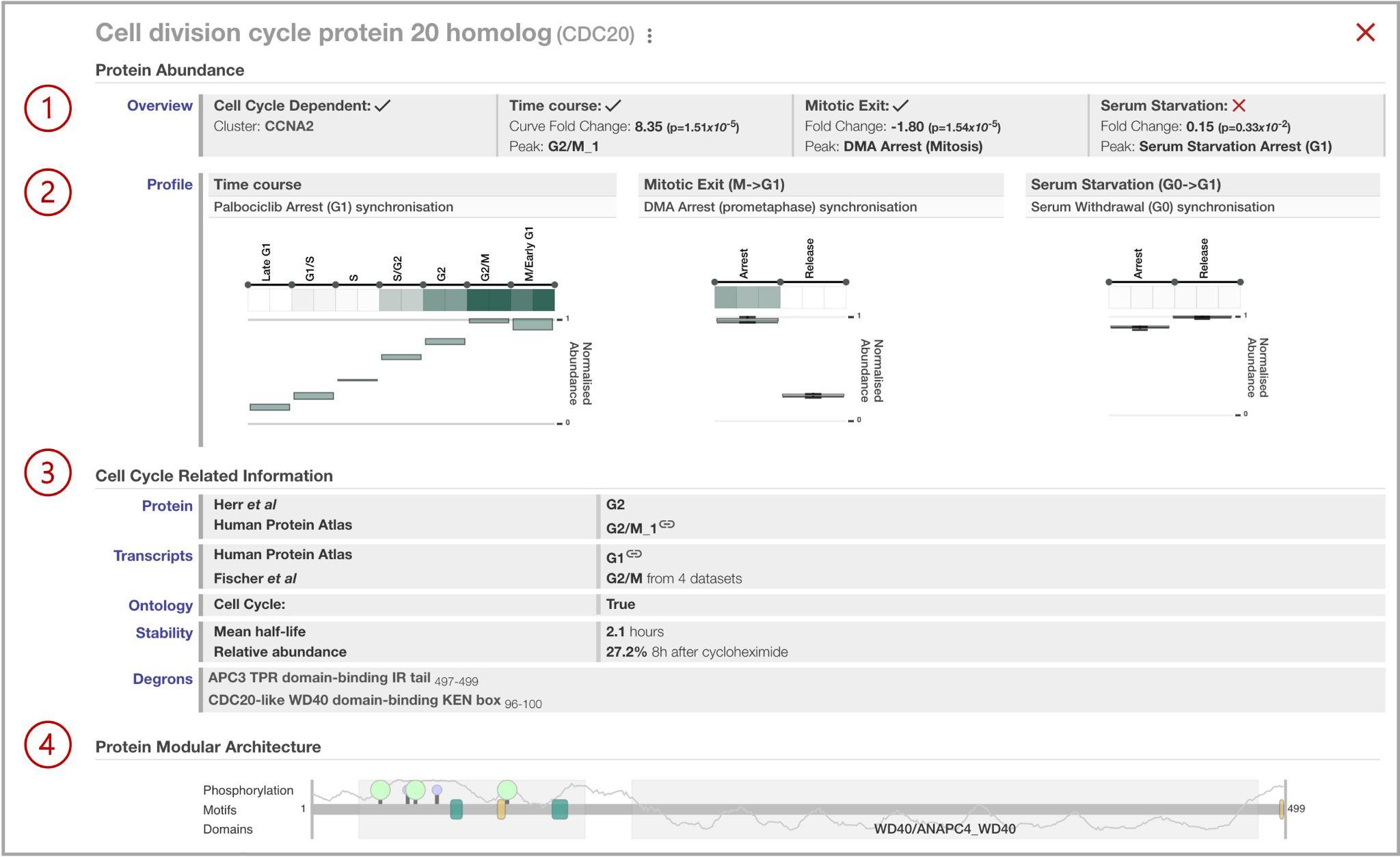

***Figure 7.*** *Overview of the Protein View page. (1) protein abundance overview section, (2) protein abundance profile section, (3) cell cycle-related information section, and (4) protein modular architecture section.*

##### ***Protein Abundance Profile Section***

The *protein abundance profile* section shows the raw data from the analysis for each dataset. In Figure 7 - (2) the log2-mean normalised protein abundances for the two Time Course replicates and three Mitotic Exit and Serum Starvation replicates are visualised as heatmaps. Below this, the 0-max protein abundance for the two Time Course replicates and three Mitotic Exit and Serum Starvation replicates is visualised as boxplots. Details of the abundance values are shown by hovering over each part of the heatmaps and the boxplots (Figure 8A).

##### ***Cell Cycle-Related Information Section***

The *cell cycle-related information* section shows relevant complementary information for external data sources (Figure 7 - 3). If there is information about the cell cycle dependency status of a protein from other publicly available proteomics and transcriptomics datasets the cell cycle phase peak coming from the respective experiment is reported. A single GO Term annotation (*cell cycle* - GO:0007049) is given to each protein to define if the protein has been annotated to this high-level ontology term or any descendants. Finally, protein stability information as defined by cycloheximide chase coupled to mass spectrometry experiments [PMID: 34626566] and annotated degrons is reported if available.

#### ***Protein Modular Architecture Section***

In the *protein modular architecture* section, (Figure 7 - 4) domain and motif features, degrons and phosphorylation sites from the CCdb screen are shown for the selected protein. All the features in this visualisation are interactive, for example, the fold change, significance and dataset are reported by hovering over the phosphorylation sites (Figure 8 B, C). Phosphorylation sites that are present in the Time Course dataset only are reported in pink, in the Mitotic Exit only in blue and if a site is present in both datasets then it is reported in green. Degrons are reported in yellow and non-degron short linear motifs (SLiMs) are reported in green. The domains are the autonomous structural units annotated based on the AlphaFold structural model [PMID: 24288371] and are represented by light grey boxes. The autonomous structural units were extracted using the xProtCAS tool [PMID:3737148] using a graph-based community detection algorithm on the AlphaFold2 predicted aligned error (PAE) matrix. Based on this data, the structural state of each residue was classified as Domain, Loop or Unstructured and autonomous structural units. The autonomous structural units can cover one or more domains and it is annotated with overlapping Pfam domains. The solvent-accessible surface windowed over 5 adjacent residues has also been calculated from AlphaFold2 models to represent the local accessibility of the protein regions. This accessibility data is represented as a grey line plot in the modular architecture. A local accessibility value indicates a more ordered the protein region. Finally, evolutionary analysis of the protein based on taxonomic range and visualised with ProViz can be accessed by clicking the ProViz button [PMID: 27085803] (Figures 8 C, D).

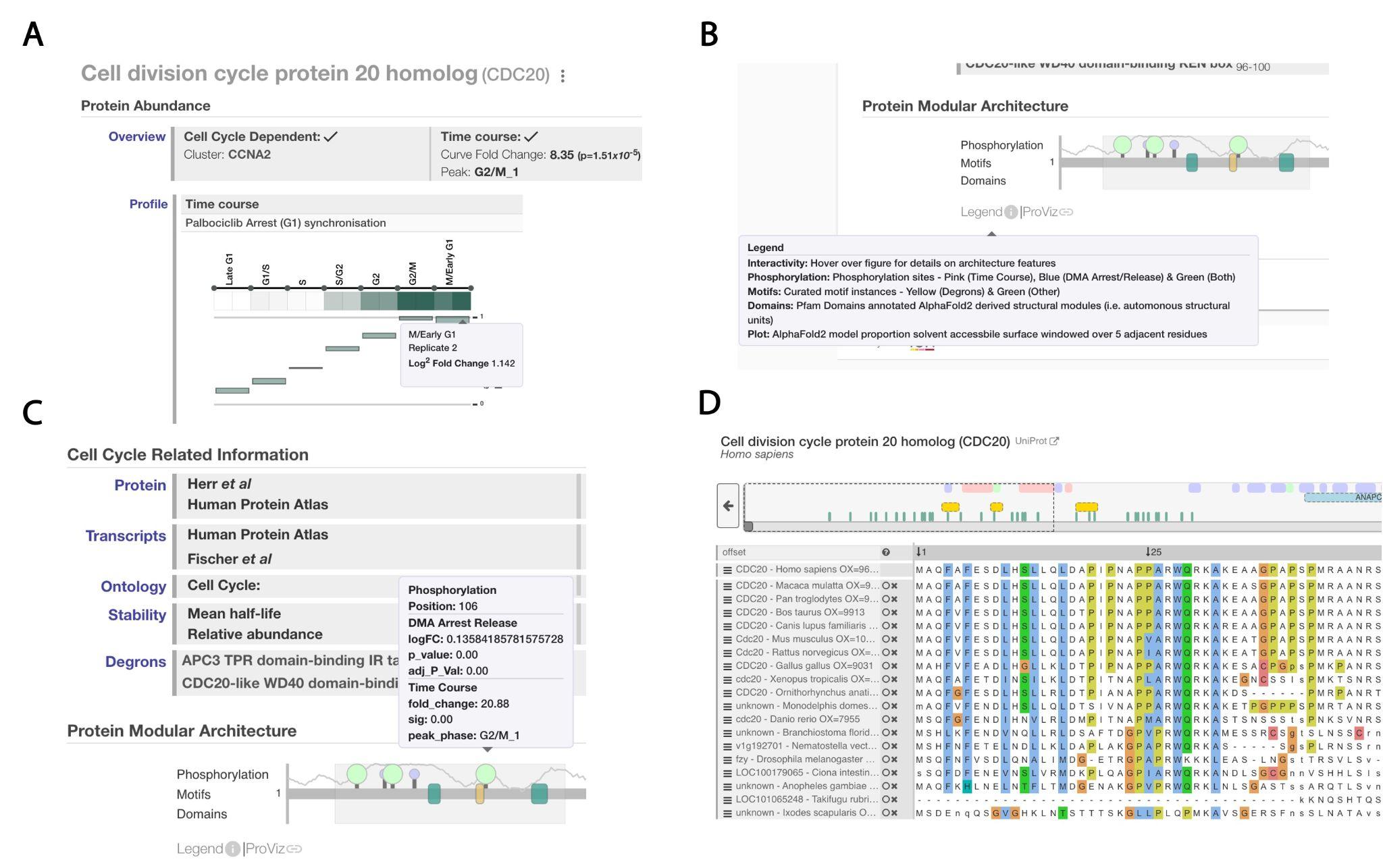

***Figure 8.*** *Overview of the interactive parts of the Protein View page showing representative examples of the tooltips that are revealed by hovering over parts of the CCdb interfaces. (A) Abundance information for the time point M/Early G1 of protein CDC20. (B) Explanation of the Protein Modular Architecture view. (C) Information for the phosphorylation site of CDC20 in position 106 of the protein. (D) Evolutionary analysis view of CDC20 provided by the ProViz tool.*

#### ***Phosphorylation Site View***

In this view, a list of all the phosphorylation events for a protein present in the CCdb dataset is provided (Figure 9). For each phosphorylation site, the normalised log2-mean abundances are shown as heatmaps for all respective replicates. The fold change and the adjusted p-value for each site in each dataset are also reported. The normalised abundances and time points are visible by hovering over the heatmaps. The cell cycle dependency status is reported by a ✓ while hovering over the cell cycle dependency status reveals the type of oscillation (protein-dependent / protein-independent). The phosphorylation sites are annotated with the AlphaFold2 accessibility, the relative conservation compared to the flanking regions and overlap with domains or SLiMs. The presence of the phosphorylation site in other PTM datasets is reported in the Modification column. All the Annotation columns of the phosphorylation view table are interactive upon hovering.

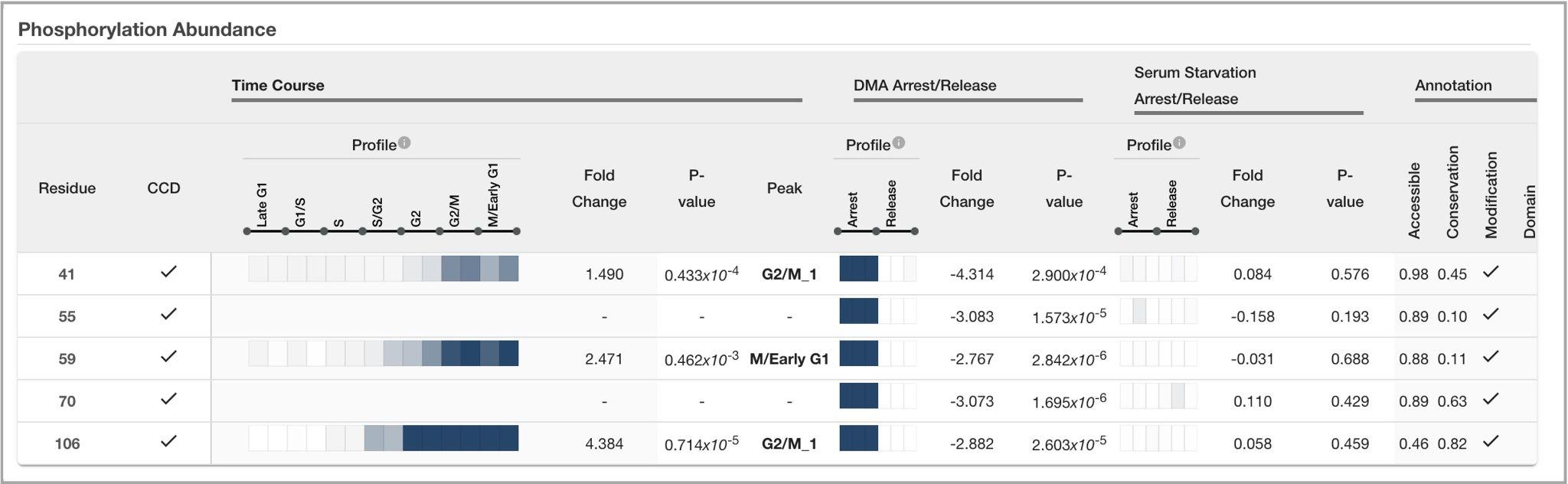

***Figure 8.*** *Example of the Phosphorylation Site View page for CDC20.*

#### ***Interactive Figures View***

The CCdb provides a resource linked to the cell cycle proteomic and phosphoproteomic data from non-cancer hTERT-immortalised retinal pigment epithelial (RPE-1) cells. This *Interactive Figures View* contains the interactive figures from the paper. In this section of the website, the user can explore the in-house datasets in-depth. All of the interactive figures can be accessed from the *Interactive Figures* tab (Figure 6-2) and there are individual legends for each figure. By hovering over dots or lines in each plot, the users can see the associated protein or phosphorylation site and the respective statistical metrics. Proteins and phosphorylation sites are also searchable for some of the plots (Figure 10 A). The functional analysis of the study can also be accessed and explored in this section. Specifically, the Gene Ontology Term Enrichment analysis, the Protein Complexes analysis, the Localisation Enrichment analysis and the DepMap Cancer Dependency annotation (Figure 10 C). Finally, this section includes heatmaps of the oscillating proteins showing the similarity of their abundance profiles to known cell cycle markers using minimum-maximum normalised Time Course dataset labelled by cell cycle phase. The clusters are grouped by colour and their respective reference protein is reported on the right. The right column shows the protein's fitted curve fold change (FC) and the Prometaphase/Early G1 time points from the Mitotic Exit dataset (Figure 10 B). Individual proteins are shown by hovering over each part of the heatmap. Additionally, each cluster can be visualised individually.

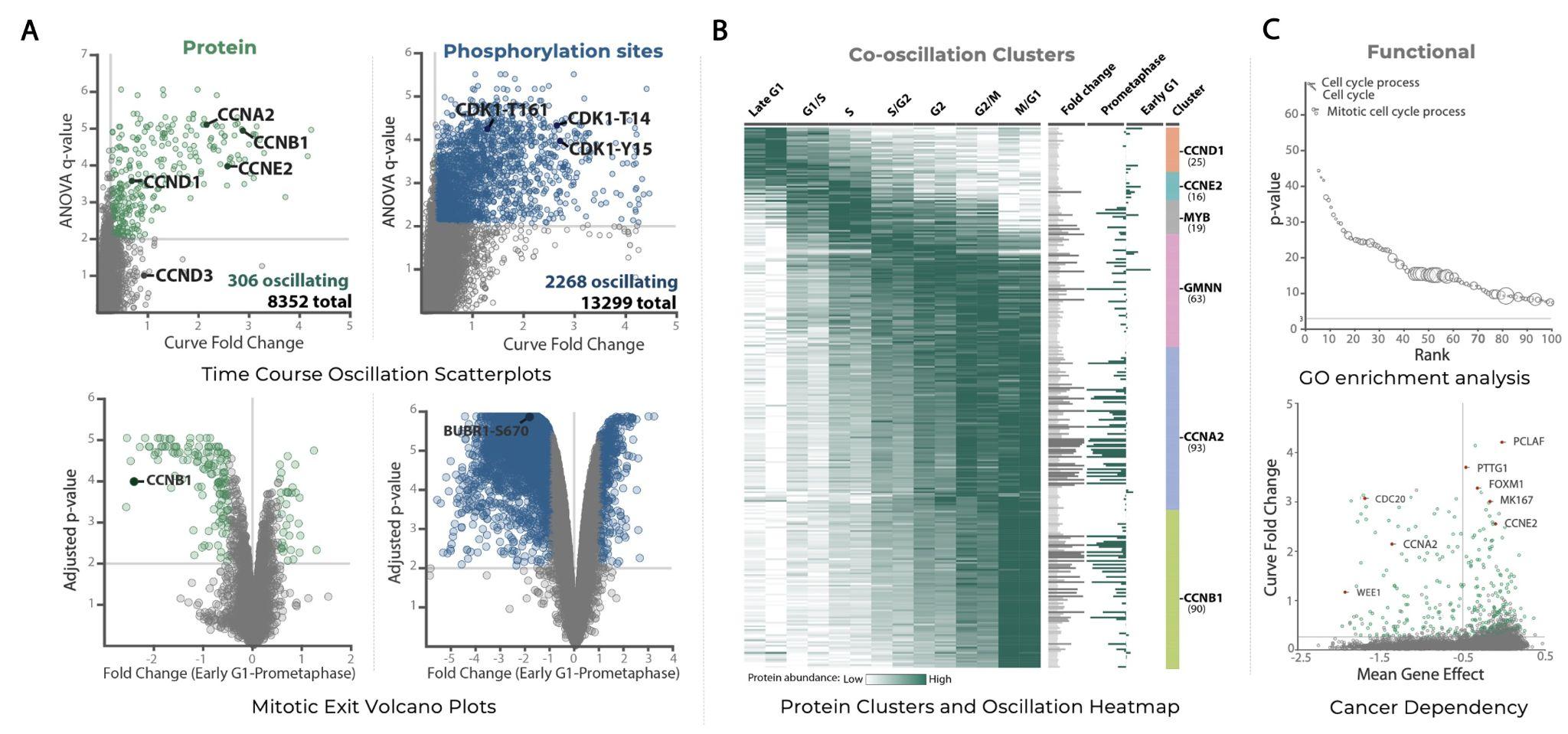

***Figure 10.*** *Examples of the Interactive Figures View page. (A) Overview of oscillation in the Time Course and Mitotic Exit datasets. (B) Co-oscillation heatmap of CCD proteins. (D) Examples of protein functional annotation.*
